# Type-B response regulators MaRR_B9 and MaRR_B12 coordinate cytokinin-mediated negative regulation of anthocyanin biosynthesis in banana fruits

**DOI:** 10.1101/2024.08.21.608915

**Authors:** Ruchika Rajput, Shivi Tyagi, Kumar Anchal, Samar Singh, Ashverya Laxmi, Prashant Misra, Ashutosh Pandey

## Abstract

- Anthocyanins are pigments responsible for vibrant plant colors and play vital roles in plant physiology. Comparing two banana varieties, grand naine (GN) and red banana (RB), which display significant contrast in anthocyanin pigmentation.
- Transcriptomic analysis of GN and RB peel and pulp tissue highlighted cytokinin-specific response regulators, MaRR_B9 and MaRR_B12, as regulators of anthocyanin biosynthesis.
- Through a combination of physiological, molecular, and biochemical analyses, we demonstrate that MaRR_B9 and MaRR_B12 exert direct regulatory control over key structural genes of anthocyanin biosynthesis, *dihydroflavanol reductase* (*MaDFR*) and *anthocyanidin synthase* (*MaANS*). MaRR_B9 and MaRRB_12 interact with the promoters of *MaDFR* and *MaANS*, repressing the concerned genes *in vivo*. Overexpression of *MaRR_B9* and *MaRR_B12* in banana fruits leads to a reduction in anthocyanin content, particularly that of the cyanidin derivative, along with altered expression patterns of *MaDFR* and *MaANS*.
- Thus, the present study identifies novel regulators of anthocyanin biosynthesis in banana and provides further evidence that the cytokinin regulatory network modifies anthocyanin accumulation in plants. In conclusion, our findings reveal new molecular targets, in the form of MaRRs for the genetic manipulation of anthocyanin content and thereby for the development of stress-resilient and nutritionally enriched crop plants.

## Introduction

Flavonoids constitute a group of specialized plant metabolites synthesized through the phenylpropanoid pathway. These compounds play diverse functions in plant biology, including their protective role against various biotic and abiotic stresses, and thereby potentially being involved in plant adaptive responses to climate change (Winkel-Shirley, 2001; Liu *et al*., 2018). Flavonoids are primarily categorized into three types based on their chemical aglycon structure: Flavonol, Anthocyanin, and Proanthocyanidin (PA). Anthocyanins, an essential pigment within this classification, are responsible for the coloration of fruits, flowers, leaves, and other plant tissues, facilitating plant reproduction through pollination and seed dispersal (Stintzing & Carle, 2004; Lev-Yadun & Gould, 2009; Khoo *et al*., 2017). Additionally, these compounds have been implicated in the absorption of excess photons, and therefore have potential to protect plants from light stress (Chalker-Scott, 1999; Tattini *et al*., 2014; Naik *et al*., 2022). In humans, anthocyanin derivatives exhibit various pharmacological properties, including anti-cancerous, anti-inflammatory, and anti-apoptotic activities. Owing to these bioactivities; anthocyanin-enriched foods offer several health benefits and are sources of nutraceutical development (Mattioli *et al*., 2020).

In the higher plants, the pathway leading to the anthocyanin biosynthesis is well-conserved and belongs to the late enzymatic steps of flavonoid biosynthesis. Dihydroxyflavonol 4-reductase (DFR) serves as a rate-limiting enzyme in anthocyanin biosynthesis, catalyzing conversion of three dihydroflavonols into colorless leucoanthocyanidin, which are subsequently transformed into colored anthocyanidins (such as cyanidin, pelargonidin, and delphinidin) by enzymes anthocyanidin synthase/leucoanthocyanidin dioxygenase (ANS/LDOX) (Liu *et al*., 2018). Anthocyanidins, initially present in an unstable aglycone form, undergo glycosylation by various glycosyltransferase enzymes, such as flavonoid 3-O-glucosyltransferase (UFGT), to form stable glycone forms of anthocyanins (Honda & Saito, 2002; Tanaka *et al*., 2008). A functionally conserved complex of three regulatory proteins belonging to MYB(M), bHLH(B) and WD40(W) families (MBW) play a pivotal role in the transcriptional regulation of genes committed to flavonoid biosynthesis in diverse plant species (Xu *et al*., 2014, 2017; Rajput *et al*., 2022a,b). The components of MBW complex specific to anthocyanin biosynthesis have been identified and characterized from diverse plant species. For example, this complex includes transcription factors, namely PAP1, PAP2, and AtEGL3, AtGL3 in Arabidopsis, and MdMYB10, and MdMYB110a, as well as MdbHLH3 and MdbHLH33 in apples (Espley *et al*., 2007; Chagné *et al*., 2013). Several exogenous and endogenous factors including light, temperature, and phytohormones modulate anthocyanin biosynthesis through modifying the expression and activity of the components of MBW complex (Naik *et al*., 2022). However, these factors may regulate anthocyanin biosynthesis independently of MBW complex by involving transcription factors from other families. For example, light signaling components, such as basic leucine zipper (bZIP) TF HY5 and PHYTOCHROMEINTERACTING FACTOR 3 (PIF3), also activate the anthocyanin pathway by directly binding to the promoters of AN-biosynthesis genes like DFR and LDOX (Shin *et al*., 2007; An *et al*., 2017). Given that multiple factors can modify anthocyanin biosynthesis, it is tempting to speculate that other unidentified transcription factors might also be involved in the transcriptional regulation of anthocyanin biosynthesis in plants. Identification of such regulators can offer novel molecular targets for the genetic manipulation of anthocyanin biosynthesis in plants.

The plant hormone cytokinin, a crucial growth regulator, induces anthocyanin biosynthesis in plants like Arabidopsis, artichoke, carrot, garden balsam, rape, and rose (Das *et al*., 2012; Wang *et al*., 2018). Recent findings by Zhu *et al*., (2023) reveal that CK induces anthocyanin biosynthesis in Eucalyptus and several other perennial woody plants but not in Arabidopsis and tobacco. This suggests a diverse role of CK in regulating anthocyanin biosynthesis across different species (Zhu *et al*., 2023). The cytokinin signal transduction is regulated by the two-component system (TCS) that incorporates histidine receptors (HKs), histidine-containing phosphotransfer proteins (AHPs) and type-B RRs. When cytokinin binds to the histidine kinase receptor, it gets autophosphorylated and relays signal to two classes of response regulators (RR): the response-promoting type-B RRs and the response-inhibiting type-A RRs via Histidine containing phospho-transfer protein (Tsai *et al*., 2012; Kieber & Schaller, 2014; Worthen *et al*., 2019). Our understanding of role of cytokinin in regulating the anthocyanin pathway in monocots is limited. The members of TCS play a crucial role in cytokinin signaling, however, the molecular mechanism concerning the cytokinin-mediated modulation of anthocyanin biosynthesis has not been studied so far.

Bananas stand as the largest tropical and subtropical fruit crop globally with various nutritional properties (https://www.statista.com/statistics/264001/worldwideproduction-of-fruit-by-variety/). Two significant cultivars of banana, Grand Naine (GN) and red banana (RB) that differ in color, and appear to accumulate anthocyanin differently, offer an intriguing background for deciphering regulatory networks involved in the biosynthesis of anthocyanin. To this end, in the present investigation, transcriptome analysis of color-contrasting banana cultivars RB vs GN at the ripening stage was carried out. The transcriptome data analysis suggested differential expression of TCS gene family members. Further, identification and phylogenetic analysis of the cytokinin responsive TCS members was carried out in banana. Since type-B RRs play a crucial role in the initial transcriptional response of plants to cytokinin, we further focused our study to characterize these regulators. Amongst, two type-B RRs (MaRR_B9 and MaRR_B12) were functionally characterized in the context of their role in the regulation of anthocyanin biosynthesis. Our study revealed that MaRR_B9 and MaRR_B12 directly binds to the promoter regions of *MaDFR* and *MaANS* and regulate their expression. Further, transient *in*-planta expression of *MaRR_B9* and *MaRR_B12* confirmed that these regulators modify the anthocyanin biosynthesis in banana. Taken together, we presented experimental evidence that cytokinin inhibits anthocyanin biosynthesis in banana, facilitated by type-B RRs, MaRR_B9 and MaRR_B12 in fruits.

## Method and materials

### Plant materials and growth conditions

Dessert bananas of the *M. acuminata* (AAA genome) cultivar Grand Naine (GN) and Red banana (RB) were cultivated at the National Institute of Plant Genome Research (NIPGR), New Delhi, India. Tissues from fruit peel and pulp were collected for subsequent gene expression and metabolite analysis. Banana fruit slices (barely ripe) were subjected to varying concentrations (0, 10, 20, and 30 μm) of 2-isopentenyladenine (iP), and samples were collected at different time points (0, 3, 6, 12, and 24 h) for both RNA isolation and phytochemical analysis. The samples for each treatment were collected in triplicates.

For transient experiments, freshly harvested banana fruits were obtained from the field. *Nicotiana benthamiana* plants were cultivated in a growth chamber (Percival AR-41L3, Perry, IA, USA) under a 16-hour light and 8-hour dark photoperiod at 22°C.

### Total anthocyanin quantification in Grand Naine and Red banana

Total anthocyanin content was assessed using the spectrophotometric method described previously (Pal *et al*., 2022). In brief, 2.5 mL of acidic methanol (1% v/v) was added to 0.25 g of lyophilized finely ground tissue. The extract was then left to incubate overnight at 4°C in the dark. Following centrifugation at 3000 g, the supernatant was collected, and absorbance was measured at 530 and 657 nm. Anthocyanin content was calculated using the formula: (A530 - 0.25 × A657)/tissue weight.

### RNA-Seq library preparation, sequencing, and differential expression analysis

The ripe peel (PL) and pulp (PP) tissue samples from GN and RB were collected and immediately frozen in liquid nitrogen. The total RNA was extracted using Spectrum™ Plant Total RNA Kit following manufacturer’s protocol. The RNA quality and quantity was assessed on agarose gel, NanoDrop and Agilent bioanalyzer. The RNA samples were treated with DNase I to remove any genomic DNA contamination. The RNA-seq libraries were prepared with RNA samples (RNA integrity number (RIN) ≥6) for Illumina sequencing according to the manufacturer’s protocol. The prepared libraries were sequenced using a high-throughput Illumina HiSeq sequencing platform. The quality control checks were performed on the raw sequencing data using FastQC to assess sequence quality and adapter contamination. During the preprocessing step of the raw reads, AdapterRemoval-v2 (version 2.2.0) was employed to trim adaptor sequences and low-quality bases, with the following modified parameters for the analysis: --minlength 30 and --trimqualities --minquality 20. Subsequently, ribosomal RNA sequences were eliminated from the preprocessed reads by aligning them with the silva database using bowtie2 (version 2.2.9) with the -N 1 parameter. The workflow then proceeds with samtools (version 1.3.1), sambamba (version 0.6.7), and BamUtil (version 1.0.13), all of which were used with their default parameters. The high-quality cleaned reads were aligned to the Banana reference genome and gene model downloaded from Banana genome hub (https://banana-genome-hub.southgreen.fr/musa_acuminata_pahang_v2). The alignment was performed using STAR program (version 2.5.3a). The resulting alignments were utilized to estimate gene and transcript expression levels using the cufflinks program (version 2.2.1). The differential expression analysis is then conducted using the cuffdiff program from the cufflinks package. The differentially expressed genes (DEGs) were obtained for all possible pairs of combinations (GN_PL vs RB_PL, GN_PP vs RB_PP, GN_PL vs GN_PP, and RB_PL vs RB_PP) with the criteria of q-value <0.05 and log2(fold change) ≥1 or ≤-1.The venn diagram of differentially expressed genes obtained in different combinations was generated by Venny 2.1 (https://bioinfogp.cnb.csic.es/tools/venny/index.html).

### Expression profiling of anthocyanin biosynthesis genes and hormone-related genes

The expression profiling of anthocyanin biosynthesis pathway genes in the transcriptome data of PL and PP tissue samples from GN and RB was done. Additionally, to elucidate the regulatory role of various hormones in color variation between GN and RB, the expression of hormone-related genes was analyzed using transcriptome data. Heatmaps were generated to visualize the expression patterns of genes associated with ethylene (ET), abscisic acid (ABA), indole-3-acetic acid (IAA or auxin), gibberellin (GA), and cytokinin pathways. The heatmap was generated using log2 FPKM value by TBtools toolkit.

### Phytohormone quantification by LC-MS

Phytohormones were quantified from ripe peel and pulp tissues of GN and RB following a method described by (Vadassery *et al*., 2012) with minor modifications. Fifty milligrams tissue was ground in liquid nitrogen and then homogenized and extracted in methanol containing internal standards. The extracted supernatant (12,000×g for 15 mins at 4 °C) was lyophilized, reconstituted in 500 *µ*L methanol, and finally injected into UPLC C18 column (Exion LC Sciex, Framingham, MA, USA) coupled to a triple quadrupole system (QTRAP6500 +; ABSciex) using an electrospray ionization.

### Identification and *in*-silico analysis of two-component system (TCS) genes

We categorized histidine kinases (HKs), type A and type B response regulators (RRs), pseudo-response regulators (PRRs) based on their domain architecture determined by SMART domain analysis tool (Letunic *et al*., 2021) and NCBI Conserved Domain Search (Wang *et al*., 2022). The nomenclature of identified HKs and RRs was assigned as previously described (Pareek *et al*., 2006; Schaller *et al*., 2007). The phylogenetic tree of both HKs and RRs was constructed by aligning protein sequences through MUSCLE alignment program and using maximum likelihood method at 1000 bootstrap replications in MEGA X software (Kumar *et al*., 2018). To visualize the conserved motifs and domains, multiple sequence alignment of HK, type A and type B RR protein sequences was created by MultAlin at default parameters (Corpet, 1988).

### Gene expression profiling by RT-qPCR in RB and GN fruit tissues

Total RNA was extracted from banana organs following the guidelines provided by the manufacturer (Sigma Aldrich) and subjected to DNase I treatment (Thermo Fisher Scientific, Waltham, MA, USA). Subsequently, reverse transcription of the total RNA was carried out to synthesize the first strand cDNA, utilizing reagents from Thermo Fisher Scientific. For quantitative real-time PCR, a mixture containing 1 μl of diluted cDNA (equivalent to 10 ng total RNA), 5 μl SYBR Green PCR Master Mix (Applied Biosystems, Waltham, MA, USA), and 5 nM of gene-specific primers was prepared, reaching a final volume of 10 μl. The quantitative real-time PCR was executed on a 7500 Fast Real-time PCR system (Applied Biosystems), using banana *Actin1* (GenBank no. AF246288) gene as an internal control. The transcript levels of each gene were assessed using the cycle threshold (Ct) 2-^ΔΔCt^ method (Livak & Schmittgen, 2001).Three biological and three technical replicates were analyzed to ensure the reliability of the results.

### Cloning of transcriptional regulators

The full-length coding sequences (CDS; without the stop codon) of the type-B RR genes *MaRR_B9* and *MaRR_B12* were amplified using first-strand cDNA of *M. acuminata* fruit tissue and a set of primers containing Gateway *attB* sites designed based on information from the Banana Genome Hub (Table S1). The resulting amplicons were cloned into the Gateway vector pDONRzeo (Invitrogen) and transformed into *Escherichia coli* TOP10 cells. Sanger sequencing of entry clones was performed by the NIPGR sequencing core facility (New Delhi, India).

### Subcellular localization of transcriptional regulators

Entry clones containing the coding sequences of *MaRR_B9* and *MaRR_B12* was recombined by Gateway LR reaction into the binary vector pSITE-3CA encoding the C terminus of yellow fluorescent protein (YFP) under the control of the CaMV 35S promoter. The resulting plasmids were transformed into Agrobacterium strain GV3101::pMP90 (Koncz & Schell, 1986). Agrobacteria harboring each construct were resuspended along with an Agrobacterium carrying a nuclear marker (NLS-RFP) (Kumar *et al*., 2018) in freshly made infiltration medium (10 mM MgCl2, 10 mM MES-KOH, pH 5.7, 150 μM acetosyringone) and co-infiltrated into the abaxial surface of *N. benthamiana* leaves and kept at 22°C for 48 h. YFP and RFP fluorescence was observed under a Leica TCS SP5 confocal laser-scanning microscope (Leica Microsystems, Wetzlar, Germany) at 514 nm excitation, 527 nm emission and 558 nm excitation, 583 nm emission wavelength, respectively.

### Identification of putative *cis*-acting elements in *proMaANS* and *proMaDFR2*

Putative target gene promoter fragment sequences were obtained from the banana genome (https://banana-genome-hub.southgreen.fr) (Martin *et al*., 2016) and 1531-bp and 1500-bp sequences upstream of the translation start of the *proMaDFR2* and *proMaANS* genes respectively were examined for AGATT *cis-*response regulators motifs using the New PLACE software (https://www.dna.affrc.go.jp), complemented by manual curation.

### Dual-luciferase reporter assay

To assess the transactivation potential of the type B RRs, the coding sequences of *MaRR_B9* and *MaRR_B12* were recombined into the destination vector pBTdest (Baudry *et al*., 2004) to create effector constructs driven by the CaMV*pro35S* promoter. Simultaneously, the entry clones of promoters (*proMaANS-1234* and *proMaDFR2-1531*) were recombined into p635nRRF containing 35S:REN (Kumar *et al*., 2018) to create the reporter constructs. The reporters were co-infiltrated with effectors into *N. benthamiana* leaves. After 48 hours, firefly luciferase and REN activity were measured in extracts from infiltrated leaf discs using the Dual-Luciferase Reporter Assay System (Promega), following the manufacturer’s protocol. Quantification was performed with the POLARstar Omega multimode plate reader (BMG Labtech, Ortenberg, Germany), and firefly luciferase activity was normalized to REN activity.

### ChIP-qPCR assay

The ChIP-qPCR assay was performed using the ChIP Plants kit (Abcam) according to the manufacturer’s instructions (Shi *et al*., 2019). Promoter fragments containing the AGATT *cis* motif cloned in p635nRRF that was used for dual-luciferase assays, were co-infiltrated with YFP-MaRR_B9 and MaRR_B12 fusion proteins into *N. benthamiana* leaves. Three days after infiltration, equally sized leaf discs were used for ChIP. Next, 2 μL anti-GFP antibody (Catalog #ab290, Abcam) was used for chromatin precipitation in two biological replicates, and 2 μL of anti-IgG antibody (Catalog#ab48386, Abcam) was used as a negative control. qPCR was performed with 5 μL 2x SYBRGreen PCR Master mix (Applied Biosystems), 1 μL ChIP DNA, and 0.5 μL each forward and reverse primer, with 40 cycles of amplification as described above. The relative fold enrichment of the target promoter fragments was calculated using the 2^−ΔCT^ method (Livak & Schmittgen, 2001). The primers used in ChIP-qPCR are listed in Supplementary Table S1.

### Transient overexpression in immature banana fruit slices

For transient overexpression studies, the coding sequences of *MaRR_B9* and *MaRR_B12* were integrated into the vector pANIC6B (Mann *et al*., 2012) and positioned under the control of the constitutive *maize Ubiquitin1* (*ZmUBI1*) promoter. The T-DNA of pANIC6B additionally contains a *proPvUBI1-GUS plus-nosT* expression cassette to monitor successful transformation through monitoring of GUS activity. Plasmids were introduced into *Agrobacterium* strain GV3101::pMP90 (Koncz & Schell, 1986), and transient overexpression experiments were performed according to Matsumoto (Matsumoto *et al*., 2009) with minor modifications. In brief, Agrobacterium cells carrying the plasmid were resuspended in infiltration medium (one-tenth strength MS salts, one-tenth strength B5 vitamins, 20 mM MES-KOH, pH 5.7, 2% (w/v) sucrose, 1% (w/v) glucose, and 200 μM acetosyringone) and kept in the dark for 3 h. Fruit discs were immersed in the Agrobacterium suspension and vacuum-infiltrated for 15 min. Excess Agrobacterium-containing medium was wiped from fruit slices, and transformed fruit discs were cultured on cultivation medium (same as infiltration medium) for 3 d and then used for GUS staining (transformation efficiency), anthocyanidin quantification and gene expression analysis.

## Results

### Transcriptome deep sequencing analysis of anthocyanin contrasting GN and RB cultivars during ripening

GN has yellow peel color on ripening due to carotenoid accumulation (Davey *et al*., 2006; Pereira & Maraschin, 2015), but is limited in anthocyanin accumulation. However, peel of RB is enriched in anthocyanins such as cyanidin 3-O-rutinoside, peonidin 3-rutinoside, malvidin-rutinoside (Fu *et al*., 2018). These observations about the differential pigmentation in the fruits of GN and RB cultivars were further corroborated by visual inspection as well as through the estimation of total anthocyanin content (Fig. 1a, b). As expected, the total anthocyanin content was found to be the highest in the peel of ripe RB banana. To understand the mechanism underlying the color-contrasting trait in GN and RB banana cultivars, the modulation at transcriptome level was studied using RNA-Seq from ripe peel and pulp tissues. The details of read alignment summary with the banana reference genome are available in Table S2. To identify common and unique set of DEGs in different pairs of combinations, venn diagrams were prepared for each set of up- and down-regulated genes (Fig. 1c). A total of 823 genes in GN_PL vs RB_PL, 1346 in GN_PP vs RB_PP, 779 in RB_PL vs RB_PP and 1629 in GN_PL vs GN_PP were uniquely up-regulated (Fig. 1c). While total 867 genes in GN_PL vs RB_PL, 1328 in GN_PP vs RB_PP, 836 in RB_PL vs RB_PP and 1569 in GN_PL vs GN_PP were uniquely down-regulated (Fig. 1c). Further, total 3595 up- and 5041 down-regulated in GN_PL vs GN_PP; 2534 up- and 3880 down-regulated in GN_PL vs RB_PL; 2775 up- and 3312 down-regulated in GN_PP vs RB_PP; and 2683 up- and 3968 down-regulated in RB_PL vs RB_PP (Fig. 1d; File S1). These observations suggest that the differences in gene expression between the peel and pulp tissues of GN and RB may contribute to their contrasting peel colors (File S1).

**Figure 1.**
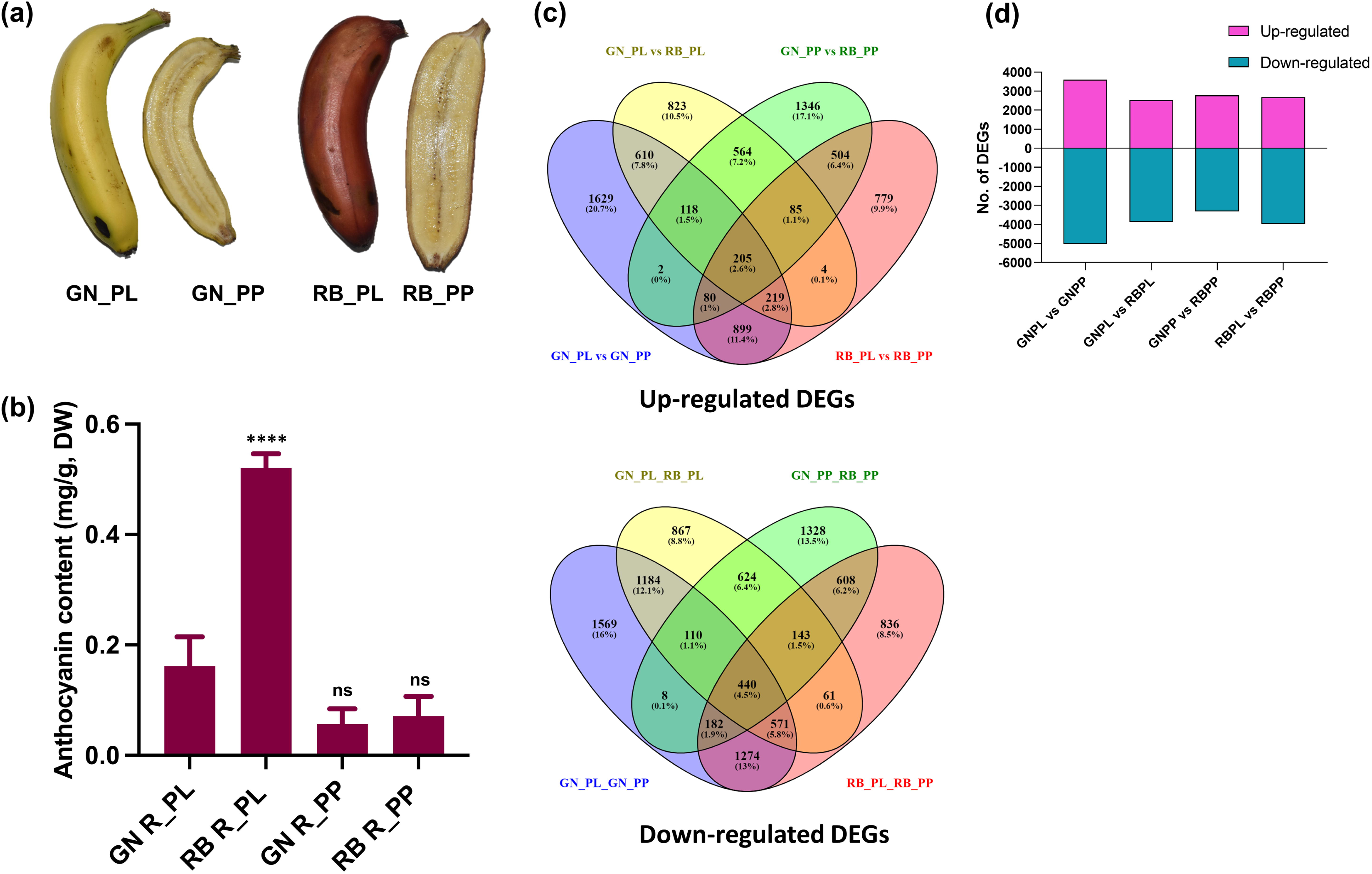
Comparative Analysis of Grand Naine (GN) and Red Banana (RB) Peel and Pulp Tissue. (a) Representative image showing the visual difference in peel color between Grand Naine (GN) and Red Banana (RB) cultivars. (b) Total anthocyanin quantification in peel and pulp tissues of GN and RB cultivars illustrating the significantly higher levels of anthocyanins in peel tissue of RB. (c) Venn diagrams depicting the differentially up-regulated and down-regulated genes (q-value < 0.05) in peel and pulp tissues of GN and RB cultivars. The diagrams illustrate the unique and shared gene expression patterns between the different tissues and cultivars, highlighting the tissue-specific gene expression changes underlying the color variation. (d) Bar graph representing the total number of upregulated and downregulated genes in peel and pulp tissues of GN and RB cultivars.

### Pathway enrichment analysis

The GO analysis of DEGs revealed enrichment of several biological processes in different combinations. Interestingly, secondary metabolic process (GO:0019748) category was enriched in down-regulated DEGs in GN_PL vs GN_PP, GN_PL vs RB_PL and RB_PL vs RB_PP indicating their role in color contrasting trait in peel tissue (Fig. S1a). The enrichment in cellular and molecular processes was also analyzed for DEGs in each GN_PL vs GN_PP, GN_PL vs RB_PL, GN_PP vs RB_PP and RB_PL vs RB_PP, showing the diversity in both the tissues of different cultivars (File S2, Fig. S1 b-c). Likewise, the KEGG pathway enrichment analysis showed that the downregulated DEGs were highly enriched in biosynthesis of secondary metabolites in GN_PL vs RB_PL as compared to other combinations (File S3, Fig. S2 a-b). The results of enrichment analysis align well with the role of secondary metabolites (anthocyanins) in the color contrasting trait of bananas.

The MAPMAN analysis was employed to visualize and analyze the transcriptome data of GN and RB for understanding the regulation overview pathways in these two varieties of banana. The expression of genes encoding transcription factors, protein modification, hormone related genes, and different antioxidant genes involved in redox signaling were modulated in regulation overview pathway in peel and pulp tissue of GN and RB (Fig. S3 a-d). Interestingly, the expression of several phytohormone-related genes were also reported to be modulated in the two varieties of banana. Based on GO mapping, KEGG pathway and MAPMAN analysis we focused our study on secondary metabolic process by studying flavonoid biosynthesis and hormone signaling genes to gain insights into the molecular mechanisms underlying their distinct phenotypic traits.

### Modulated expression of anthocyanin biosynthesis genes in GN and RB fruit

The RB peel is rich in anthocyanin content, which contributes to its distinctive color. Therefore, we investigated the expression of anthocyanin biosynthesis genes in ripe peel and pulp tissues of GN and RB. We found that the early biosynthesis genes, including *CHS*, *CHI*, and *F3H*, showed enhanced expression in the ripe peel tissue of both cultivars compared to the pulp tissue.

Moreover, the expression of certain genes was significantly higher in GN_PL compared to RB_PL. For instance, the expression of two *CHS* genes, *Ma06_g12370* and *Ma06_g17870*, was 10 and 5 times higher, respectively. The expression of a *CHI* (*Ma11_g21950*) was 7 times higher, while the expression of two *F3H* genes, *Ma02_g04650* (*MaF3H1*) and *Ma07_g17200* (*MaF3H2*), was 4 and 43 times higher, respectively.

We also analyzed the expression of late biosynthesis genes *DFR* and *ANS*. The expression of *DFR1* (*Ma03_g32330*), *DFR2* (*Ma04_g10620*) and *ANS* was 2.7, 19 and 14 times higher, respectively, in GN_PL compared to RB_PL. However, the expression of certain *UFGTs*, such as *Ma04_g10370*, *Ma06_g06990*, and *Ma07_g25160*, was 44, 4, and 5 times higher, respectively, in RB_PL compared to GN_PL (Fig. 2a). In addition, the expression levels of other flavonoid biosynthesis genes differed between RB_PL and GN_PL. Specifically, the *FLS* (*Ma03_g06970*) showed a 1.8-fold higher expression in RB_PL than in GN_PL. Conversely, the *LAR* (*Ma05_g16480*) exhibited a 1.7-fold higher expression and the *ANR* (*Ma08_g01380*) displayed 11-fold higher expression in GN_PL compared to RB_PL (Fig. 2a). These results suggest that differential expression of genes involved in flavonoid biosynthesis might contribute to the development of distinct pigmentation phenotypes of the two varieties.

**Figure 2.**
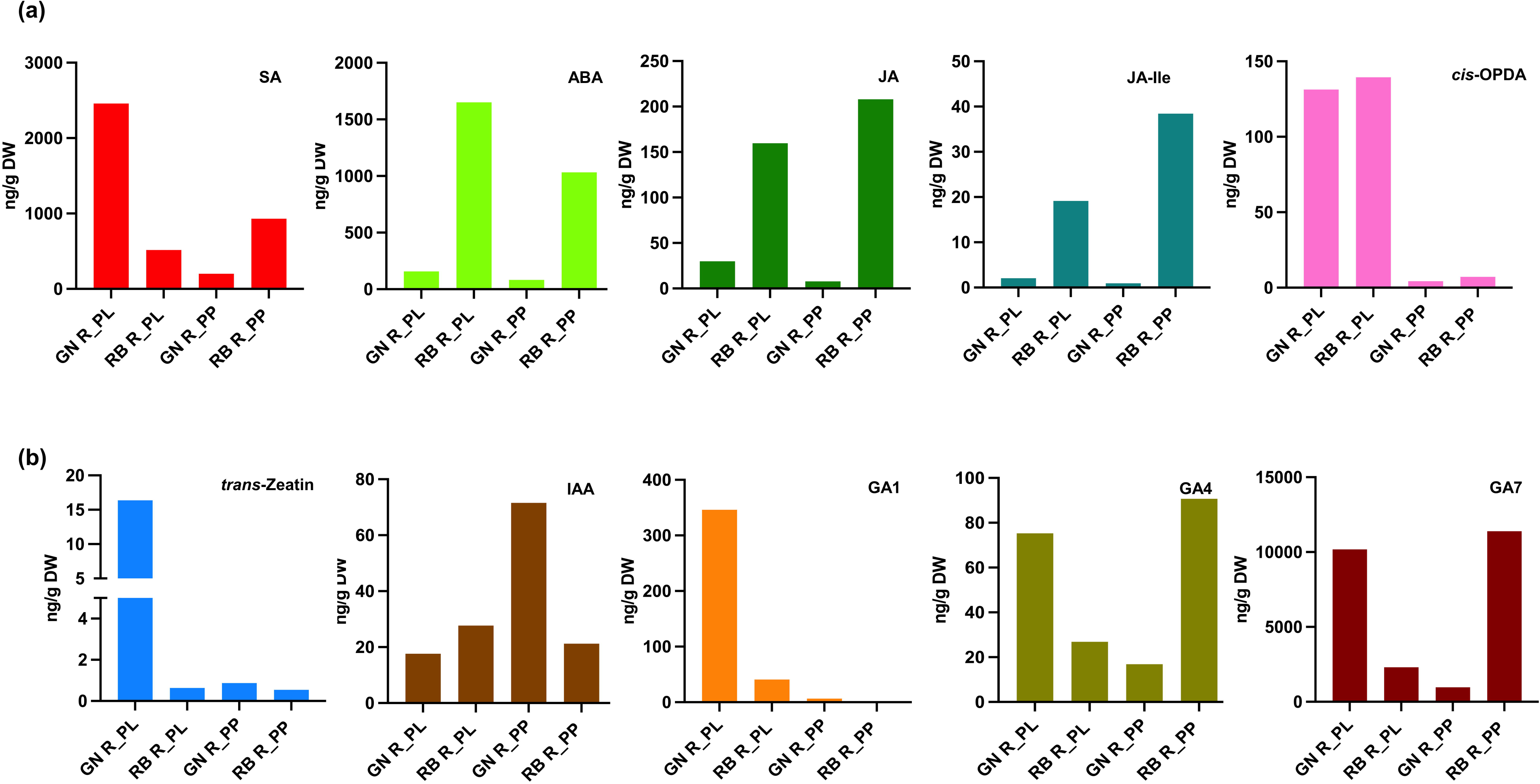
Expression profiling of anthocyanin biosynthesis genes and hormone-responsive genes. (a) Schematic representation of the anthocyanin biosynthesis pathway showing key enzymes involved in the pathway. Heatmaps illustrate the expression profiles (Log2 FPKM values) of anthocyanin biosynthesis genes in peel (PL) and pulp (PP) tissue of Grand Naine (GN) and Red Banana (RB) cultivars. (b) Heatmaps illustrating the expression profiles of ethylene (ET), abscisic acid (ABA), indole-3-acetic acid (IAA), gibberellic acid (GA), and cytokinin associated genes in GN_PL vs RB_PL, GN_PP vs RB_PP, GN_PL vs GN_PP, and RB_PL vs RB_PP. Colorbar scale from aqua-blue to purple represents low to high expression.

### Modulated expression of phytohormone-related genes in GN and RB fruit

Since phytohormones has an important role in the regulation plant growth, development, ripening and metabolic pathways, the transcriptome data was analyzed to study the modulation of genes associated with phytohormone pathways. To this end, the analysis suggested differential regulation of the genes concerning indole-3-acetic acid (IAA), abscisic acid (ABA), gibberellin (GA), ethylene (ET), and cytokinin pathways in peel and pulp tissues of GN and RB (Fig. 2b). Amongst, the transcript levels of certain genes such as ACS (*Ma05_g13700*), ERFs (*Ma03_g24970*, *Ma04_g06840*, *Ma04_g21160*, *Ma04_g21170*, *Ma04_g39770*, *Ma07_g03740*, *Ma07_g06960*,*Ma10_g19950*, and *Ma10_g26920*), GA2OX (*Ma09_g06290*), PP2Cs (*Ma03_g03910*, *Ma07_g15490*, *Ma07_g23800*) and type B-RR (*Ma10_g03650*) was >1.5 folds higher in RB_PL as compare to all other tissues. Contrary, the expression of many genes such as ACO (*Ma06_g14400*,*Ma06_g14410*), EIL (*Ma05_g30040*), ERF (*Ma06_g09750*, *Ma07_g21320*), ABA2 (*Ma07_g01480*), PP2C (*Ma09_g24350*), SnRK2 (*Ma06_g16710*), GA20OX (*Ma08_g15080*), GA2OX (*Ma04_g15130*), ASA (*Ma09_g07380*), TIR/AFB (*Ma10_g12200*), IAA/AUX (*Ma03_g03200*, *Ma09_g26000*), ARF (*Ma04_g16460*, *Ma06_g09690*, *Ma06_g14250*), type A-RR (*Ma09_g19530*), type B-RR (*Ma01_g15650*), etc. was >2 folds higher in GN_Peel than in all other tissues (Fig. 2b).The modulation in the expression levels of these phytohormones-related genes might play a crucial role in defining the phenotypes of the two varieties including the differential biosynthesis of anthocyanin.

The difference in gene expression between GN and RB peel can be attributed to a variety of factors, including differences in anthocyanin content, ripening processes, fruit development, etc. Since the role of two-component system (TCS) genes mediating cytokinin signal transduction is unexplored till date. Only a few previous findings have indicated the role of cytokinins in anthocyanin biosynthesis (Deikman & Hammer, 1995; Das *et al*., 2012). Here, we were interested in understanding the genetic regulation of anthocyanin biosynthesis by cytokinin pathway TCS genes.

### Quantitative estimation of phytohormones involved in the growth and defense

We conducted targeted phytohormone profiling using LC-MS to quantitatively estimate individual hormones. Three defense hormones, Salicylic acid (SA), Abscisic acid (ABA), and Jasmonic acid (JA, including JA-Ile and *cis*-OPDA), were quantified (Fig. 3a), along with three growth hormones, namely cytokinin (*trans*-zeatin), auxin (IAA), and the major bioactive gibberellic acids (GA1, GA4, GA7) (Fig. 3b). SA predominantly accumulated in GN_PL and RB_PP, while ABA and JA precursors were mainly present in RB_PL and RB_PP tissues. Whereas growth hormones such as *trans*-zeatin, IAA, and GAs were detected in GN_PL, RB_PP, and GN_PP tissues. The differential accumulation of phytohormones again suggests their important role in the varying anthocyanin accumulation in the two cultivars. Interestingly, the amount of *trans*-zeatin is significantly higher in GN_PL tissue which further increase our interest in understanding the genetic regulation of anthocyanin biosynthesis by cytokinin pathway.

**Figure 3.**
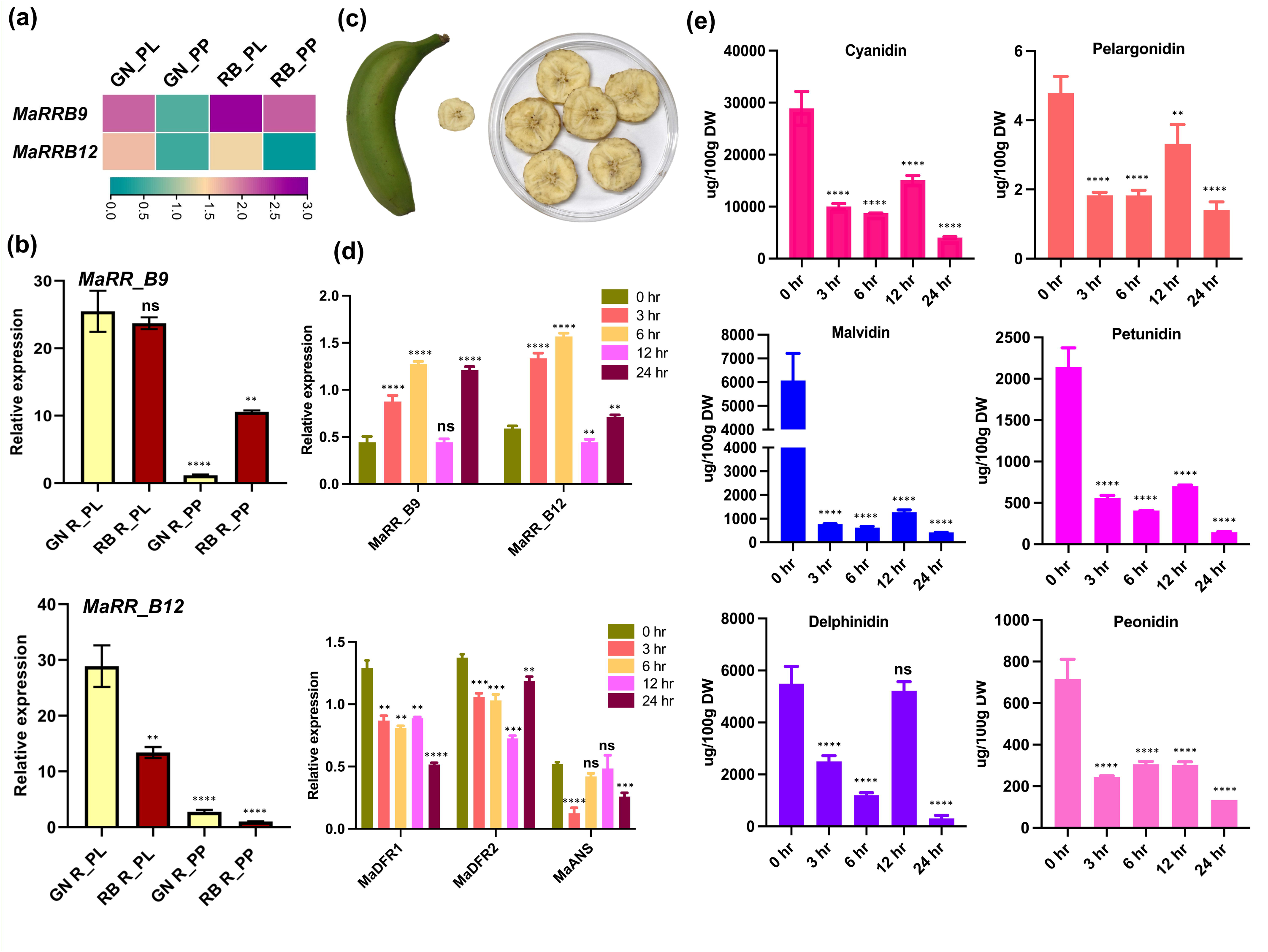
Amount of Phytohormones involved in the growth and defense response determined by LC-MS: Estimation of (a) Defence and (b) Growth hormones in ripe peel (PL) and pulp (PP) tissue of Grand Naine (GN) and Red Banana (RB). SA, Salicylic acid; ABA, Abscisic acid; JA, Jasmonic acid; JA-Ile, jasmonoyl isoleucine; *cis*-OPDA,12-oxophytodienoic acid; IAA, indole-3-acetic acid; GA, Gibberellic acid.

### Identification and phylogenetic analysis of cytokinin pathway two component system (TCS) proteins from *M. acuminata*

Based on the analysis of differentially expressed genes concerning phytohormone pathway, it was assumed that phytohormone signaling might contribute to the differential accumulation of anthocyanin. Since very limited knowledge is available on the signaling mechanism associated with cytokinin mediated regulation of anthocyanin, we focused on the genes involved in the cytokinin signaling. To this end, firstly, a detailed study on the TCS genes mediating cytokinin signal transduction was carried out using banana genome.

The cytokinin pathway TCS proteins were distinguished by their domain structure. Nineteen protein sequences containing a phosphoacceptor receiver (REC) domain were categorized as type A-RR, while twelve sequences with a REC domain at the N-terminal and a MYB-like DNA binding domain known as GARP (GOLDEN/ARR/Psr1) at the C-terminal were designated as type B-RR. Moreover, ten members of a distinct subgroup of RRs, known as Pseudo RRs (PRRs) were identified, which do not possess the conserved aspartate residues necessary for phosphorylation (Tiwari *et al*., 2021). Further, a total of 11 cytokinin HKs, which consisted of a cyclase/HK-associated sensory extracellular (CHASE) domain, histidine-kinase (HK) domain and a receiver (Rec) domain, were identified. Additionally, all HKs except *Ma01_g16190* comprises a Histidine kinase-like ATPases (HATPase_C) domain. Nine protein sequences were identified as histidine phosphotransfer (HpT) proteins, which comprised of conserved histidine phosphotransfer domain, and are responsible for mediating phosphotransfer reactions.

We separately performed the phylogenetic analysis of RRs, HKs, and HpTs from *M. acuminata* with already known sequences of these classes from *Arabidopsis* and *O. sativa*. The phylogenetic tree of RRs showed clustering into three major groups i.e., type A-RR, type B-RR and PRR (Fig. 4a). However, a few proteins from *Arabidopsis* and *O. sativa* were outlier due to their sequence diversity and presence of some addition domains. Most of the MaRRs depicted tight clustering with the OsRRs instead of ARRs, indicating divergence between monocots and dicots. Multiple sequence analysis of type B-RRs of *M. acuminata* revealed high conservation in REC domain and MYB-like DNA binding domain (Fig. 4b, Fig. S4). The phylogenetic analysis and multiple sequence alignment of each HKs (Fig. S5 a-b) and HpTs (Fig S6 a-b) showed formation of three tightly clustered groups and conserved domain architecture revealing the evolutionary relationships among the analyzed sequences.

**Figure 4.**
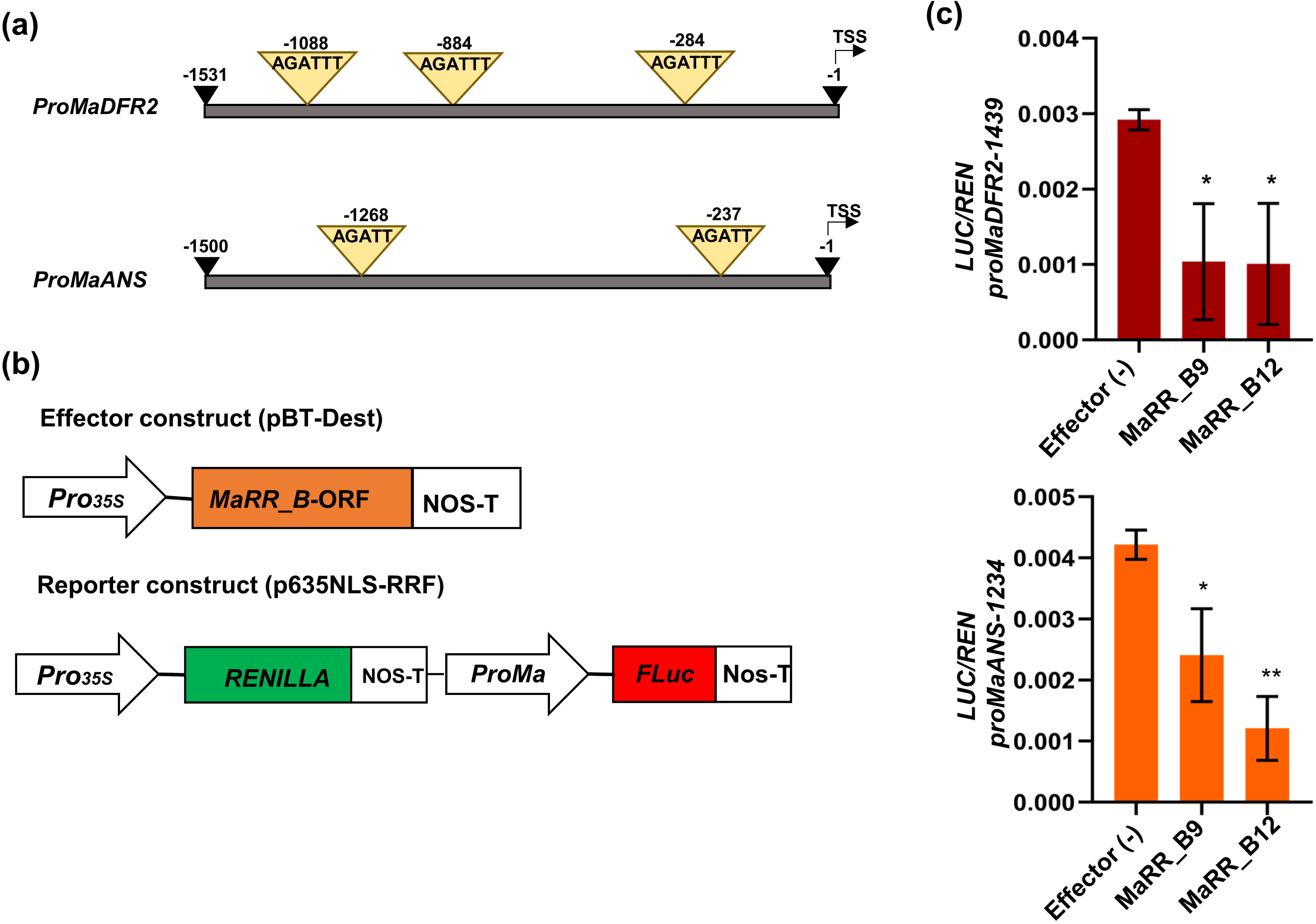
Phylogenetic tree, multiple sequence alignment, and subcellular localization. (a) Phylogenetic tree illustrating the evolutionary relationships among response regulators (RRs) from *Arabidopsis thaliana*, *Oryza sativa*, and *Musa acuminata*. RRs are grouped into type-A RRs (shaded in blue), type-B RRs (in green), and pseudo-RRs (in pink), with some members identified as outliers. The tree was constructed using the maximum-likelihood method with 1000 bootstrap replicates. Bootstrap values are denoted by brown circles at the nodes, with circle size indicating bootstrap values and branch length on each branch. (b) Multiple sequence alignment of the receiver (Rec) domain and MYB-like DNA binding domain of type-B RRs from banana, Arabidopsis, and rice, highlighting conserved amino acid residues marked with black stars. (c) Subcellular localization of 35S::MaRR_B9::YFP and 35S::MaRR_B12::YFP fusion proteins analyzed by confocal microscopy in Agrobacteria-infiltrated *Nicotiana benthamiana* leaves, demonstrating fluorescence in the nucleus. Fluorescence from the NLS nuclear RFP marker is also visible in the nucleus. Superimposed images of 35S::MaRR_B9::YFP and 35S::MaRR_B12::YFP fusion proteins with the NLS nuclear RFP marker confirm the nuclear localization of MaRR_B9 and MaRR_B12 proteins.

### Real time expression profiling of TCS genes in fruit tissues of banana

Following *in-silico* expression profiling of cytokinin-associated genes, we identified *MaTCS* genes that exhibited differential expression in GN_PL vs RB_PL and GN_PP vs RB_PP and subsequently their real-time expression analysis was conducted. The variance in expression levels of TCS genes in the peel and pulp tissues of RB and GN signifies their possible involvement in the anthocyanin biosynthesis. The expression of most TCS genes including MaHKs (*Ma04_g16280*, *Ma04_g18090*, and *Ma11_g13610*), MaRRAs (*Ma01_g21240* and *Ma03_g08570)*, and MaRRBs (*MaRR_B5*, *MaRR_B8*, *MaRR_B11* and *MaRR_B12*) displayed higher expression in GN_PL tissue compared to GN_PP, RB_PL and RB_PP tissues. Notably, certain genes such as *Ma08_g16680*, *MaRR_B6*, and *MaRR_B9* has higher expression levels in GN_PL and RB_PL, while RB_PP and GN_PP has almost similar expression patterns of these genes (Fig. 5a, 5b, Fig. S7, S8). Overall, the expression profiling validated the differential expression of TCS genes, reported by transcriptome analysis. The differential regulation of TCS genes might play an important role in defining differential accumulation of anthocyanin in the two pathways. We focused here on two homologous type-B RRs *MaRR_B9* (Ma10_g03650) and *MaRR_B12* (Ma11_g07910) as potential candidates for further characterization because *MaRR_B12* is differentially expressed in peel tissue of GN and RB (the major site for AN biosynthesis), as indicated by both the transcriptome and RT-qPCR analyses. MaRR_B9, on the other hand shows close homology with MaRR_B12 so we also selected it for functional characterization.

**Figure 5.**
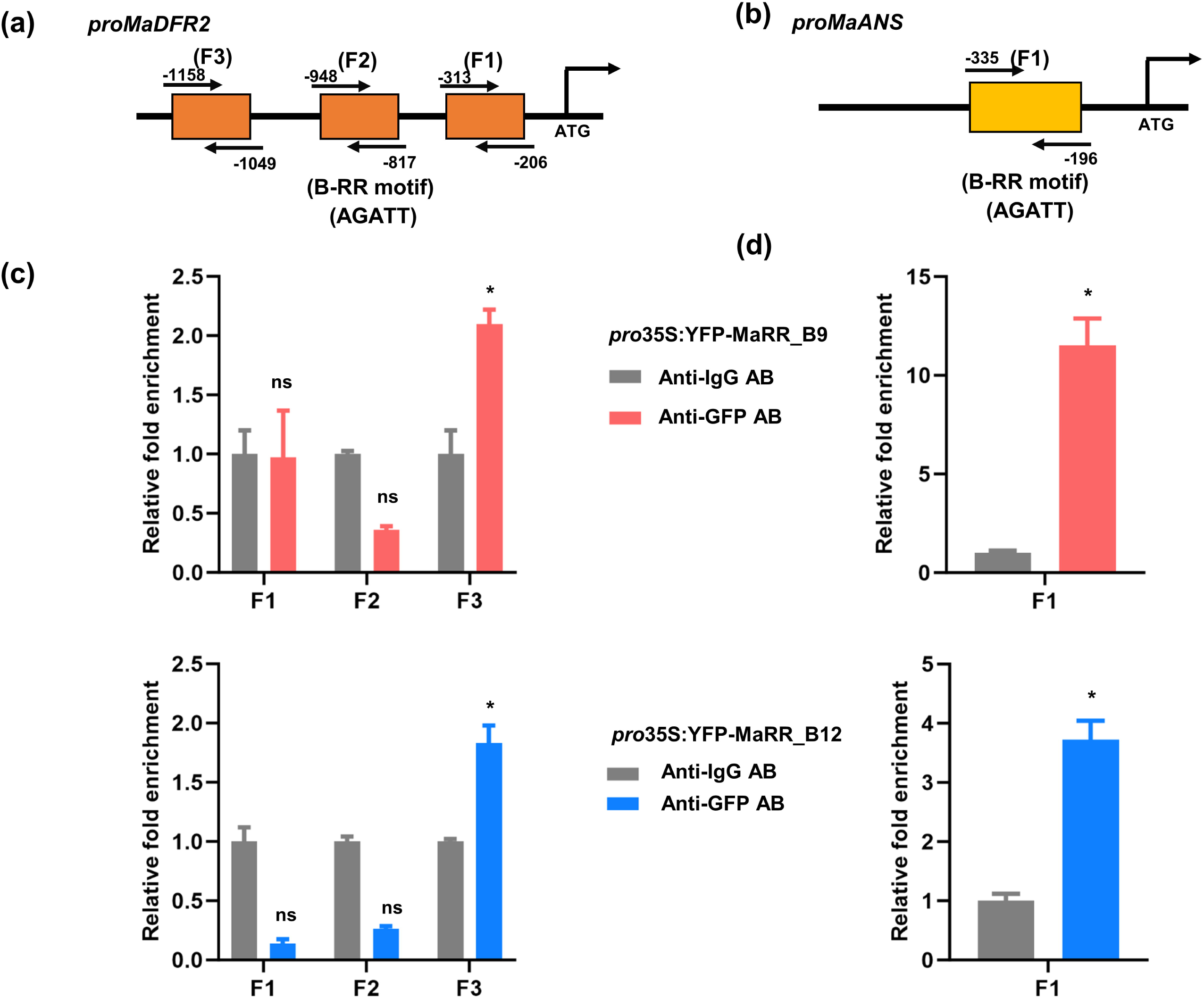
Expression analysis and anthocyanin quantification in cytokinin-treated banana fruit. (a) Heatmap showing the expression of *MaRR_B9* and *MaRR_B12* in peel (PL) and pulp (PP) of GN and RB using log2(FPKM) values. Color bar scale from aqua-blue to purple represents low to high expression (b) RT-qPCR expression analysis of *MaRR_B9* and *MaRR_B12* in PL and PP tissues of GN and RB. (c) Representative image showing the banana (barely ripe) and its slices used for cytokinin treatment with different 2-isopentenyladenine (2-iP) concentrations (0, 10, 20, 30 µM). (d) Expression profiling of *MaRR_B9*, *MaRR_B12*, *MaDFR1*, *MaDFR2*, and *MaANS* genes at different time points (0, 3, 6, 12, and 24 h) of 30 μM 2-ip treatment by RT-qPCR. (e) Bar graphs represent anthocyanins (cyanidin, malvidin, delphinidin, pelargonidin, petunidin, and peonidin) content in μg/100g dry weight (DW) at different time points (0, 3, 6, 12, and 24 h) of 30 μM 2-ip treatment. The graph shows values ± SD of two independent biological replicates each having three technical replicates and the error bars give ±SD values. *MaActin1* expression was used as reference control for RT-qPCR.

We also performed the subcellular localization of MaRR_B9 and MaRR_B12 by transiently co-infiltrating constructs encoding a YFP fusion in *N. benthamiana* epidermal cells and the nuclear marker NLS-RFP (red fluorescent protein (RFP) fused to a nuclear localization signal (NLS)). We detected YFP fluorescence exclusively in the nucleus with co-localized NLS-RFP indicating that MaRR_B9 and MaRR_B12 are nuclear proteins (Fig. 4c).

### Cytokinin treatment lowers the expression of anthocyanin biosynthesis genes and the content of anthocyanin derivatives in banana fruit tissues

The gene expression analysis suggested possible role of cytokinin signaling in modulating anthocyanin biosynthesis in the two varieties of banana. Therefore, we examined the expression of genes involved in anthocyanin biosynthesis after exposure of varying concentrations of 2-isopentenyladenine (iP) [0, 10, 20, 30 µM] to barely ripe 18 w old fruit pulp tissues followed by systematic collection and analysis of samples at varying time points: 0, 3, 6, 12, and 24 h (Fig. 5c). The RT-qPCR data revealed that the treatment with 30 µM iP significantly lowers the expression of anthocyanin biosynthesis genes *DFR1, DFR2* and *ANS* (Fig. 5d). Interestingly, the expression of *MaRR_B9* and *MaRR_B12* was increased followed by cytokinin treatment in pulp tissues (Fig. 5d). Further, metabolite analysis also demonstrated a decrease in the content of anthocyanin derivatives in cytokinin treated fruit samples following 2-iP treatment at 30 µM (Fig 5e).The collective gene expression and metabolite data strongly suggested that the exogenous 2-iP treatment leads to an increase in the transcriptional level of *MaRR_B9* and *MaRR_B12*, and decrease in the expression of genes associated with anthocyanin biosynthesis, consequently lowering the anthocyanin content in banana fruit tissues.

### Type B RRs binds to the *MaDFR* and *MaANS* promoters and downregulates their expression

Furthermore, the promoters of *MaDFR* and *MaANS* were retrieved and studied for the presence of various *cis*-regulatory elements. After detailed structural analysis, the B-ARRs motif AGATT (Xie *et al*., 2018) was identified in the promoter regions of *MaDFR* and *MaANS,* suggesting that MaRRs might directly regulate the transcription of *MaDFR2* and *MaANS*. Within the 1.5 kb *proMaDFR2* fragment, three binding sites were identified, paralleled by two corresponding binding sites in the 1.5 kb region of the *proMaANS* (Fig. 6a). Additionally, one putative binding site was identified in the promoter fragment of *proMaDFR1* (Fig. S9). In terms of quantity and distance, we found an accumulation of maximum putative binding sites in the selected regions of the*MaDFR2* and *MaANS* promoters, and therefore, we decided to use these promoter fragments for further analyses. To test the transactivation potential of *MaRR_B9* and *MaRR_B12* on the promoters of their putative target genes pro*MaDFR2-1531* and pro*MaANS*-*1234*, Agroinfiltration-based transient assay followed by dual luciferase assay was carried out. The reporter constructs consisted of the promoters *proMaDFR2* and *proMaANS* to be analyzed was placed upstream of the *firefly luciferase* (LUC) reporter gene, with *Renilla luciferase* (REN) serving as a normalization gene (Fig. 6b). The effector constructs, containing the coding sequences of *MaRR_B9* and *MaRR_B12* were co-infiltrated alongside the reporter constructs into *N. benthamiana* leaves. The results indicated that *MaRR_B9* and *MaRR_B12* significantly reduced the transactivation potential of pro*MaDFR2* and pro*MaANS*, providing further insights into the regulatory role of type-B RRs in anthocyanin biosynthesis (Fig. 6c). It suggests that MaRR_B9 and MaRR_B12 might repress the expression of *MaDFR2* and *MaANS* by directly binding to their promoter regions.

**Figure 6.**
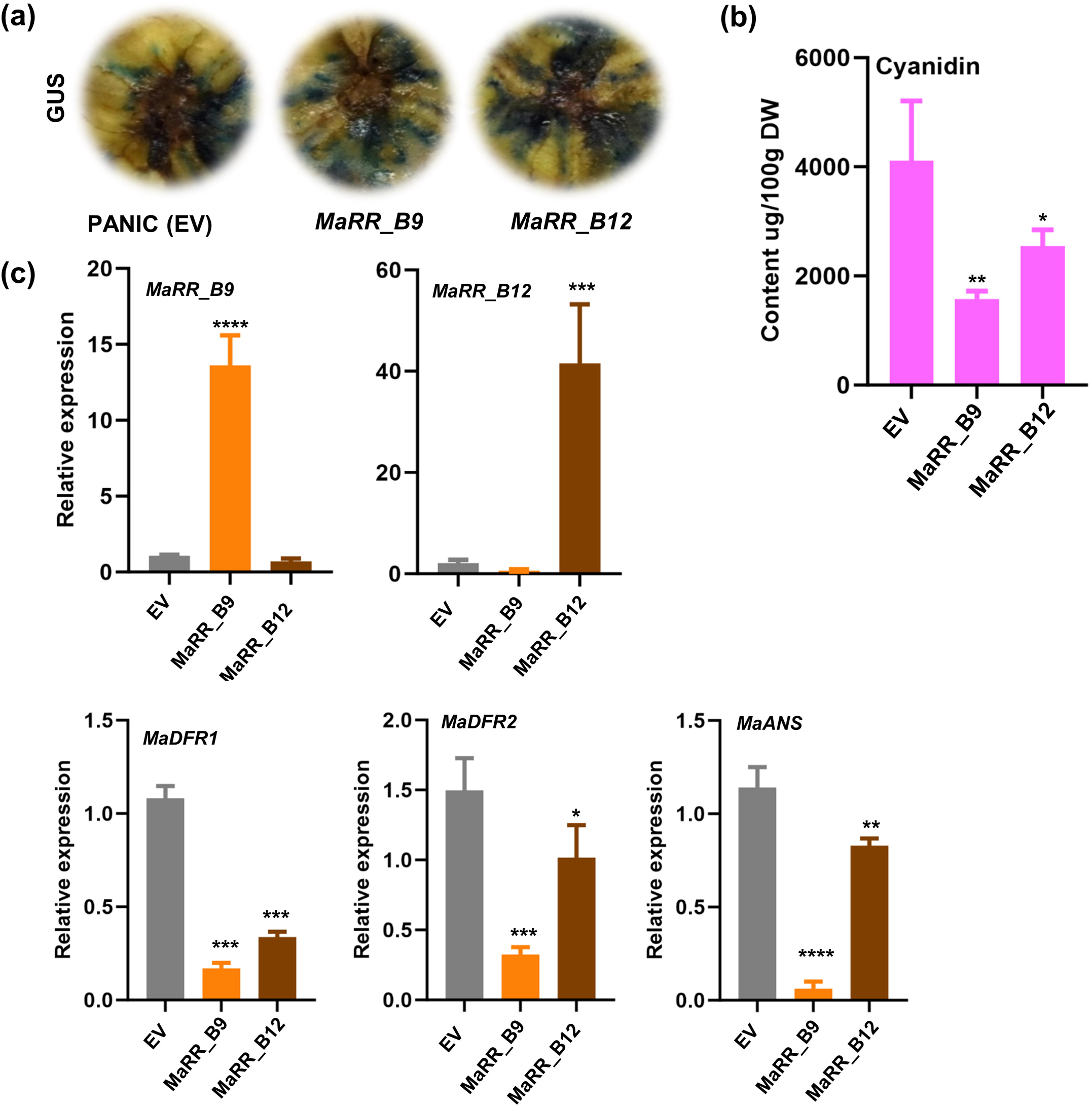
MaRR_B reduces *MaDFR2* and *MaANS* level by binding to the RR elements in their promoters: (a) Schematic diagram of the *MaANS* and *MaDFR2* promoters with putative *cis*-acting RR (AGATnn) sites as indicated with yellow triangles. The black triangles mark the − 1 position relative to the transcription start site (TSS). (b). Schematic diagram of the effector and reporter constructs used in the dual luciferase experiment (c) Results from transient luciferase assays in *Nicotiana benthamiana* leaves. Constructs harbouring the firefly luciferase reporter gene (LUC) driven by 1531-bp *MaDFR2* and 1234-bp *MaANS* promoter fragments were transiently co-infiltrated with MaRR_B9 and MaRR_B12 effector constructs. *, p ≤ 0.05; **, p ≤ 0.01, ***, p ≤ 0.001, as determined by one-way ANOVA. Data are shown as means ± SD of four biological replicates.

### *MaRR_B9* and *MaRR_B12* occupies gene promoters of *MaDFR2* and *MaANS*

Next, to verify the direct binding of MaRRs to the B-RR motifs in *proMaDFR2* and *proMaANS*, we co-infiltrated *pro35S:YFP-MaRR_B9* and *pro35S:YFP-MaRR_B12* with *proMaDFR2* and *proMaANS* constructs into *N. benthamiana* leaves. Subsequently, a chromatin immunoprecipitation (ChIP) assay was performed using an anti-GFP antibody to immunoprecipitate YFP-MaRR_B bound chromatin and anti-IgG was used as a negative control. ChIP-qPCR with purified DNA from the precipitated chromatin showed enrichment of the fragment F3 containing B-RR element of *proMaDFR2* whereas F1 and F2 did not showed any enrichment and for *proMaANS,* fragment F1 showed enrichment (Fig 7a, 7b, 7c, 7d). These results revealed that MaRRs directly bind to the F3 and F1 fragment of pro*MaDFR2 and proMaANS,* respectively and regulate their expression. These observations confirmed that MaRR_B9 and MaRR_B12 act as repressors of ANS and DFR2, and thereby regulate the anthocyanin biosynthesis.

**Figure 7.**
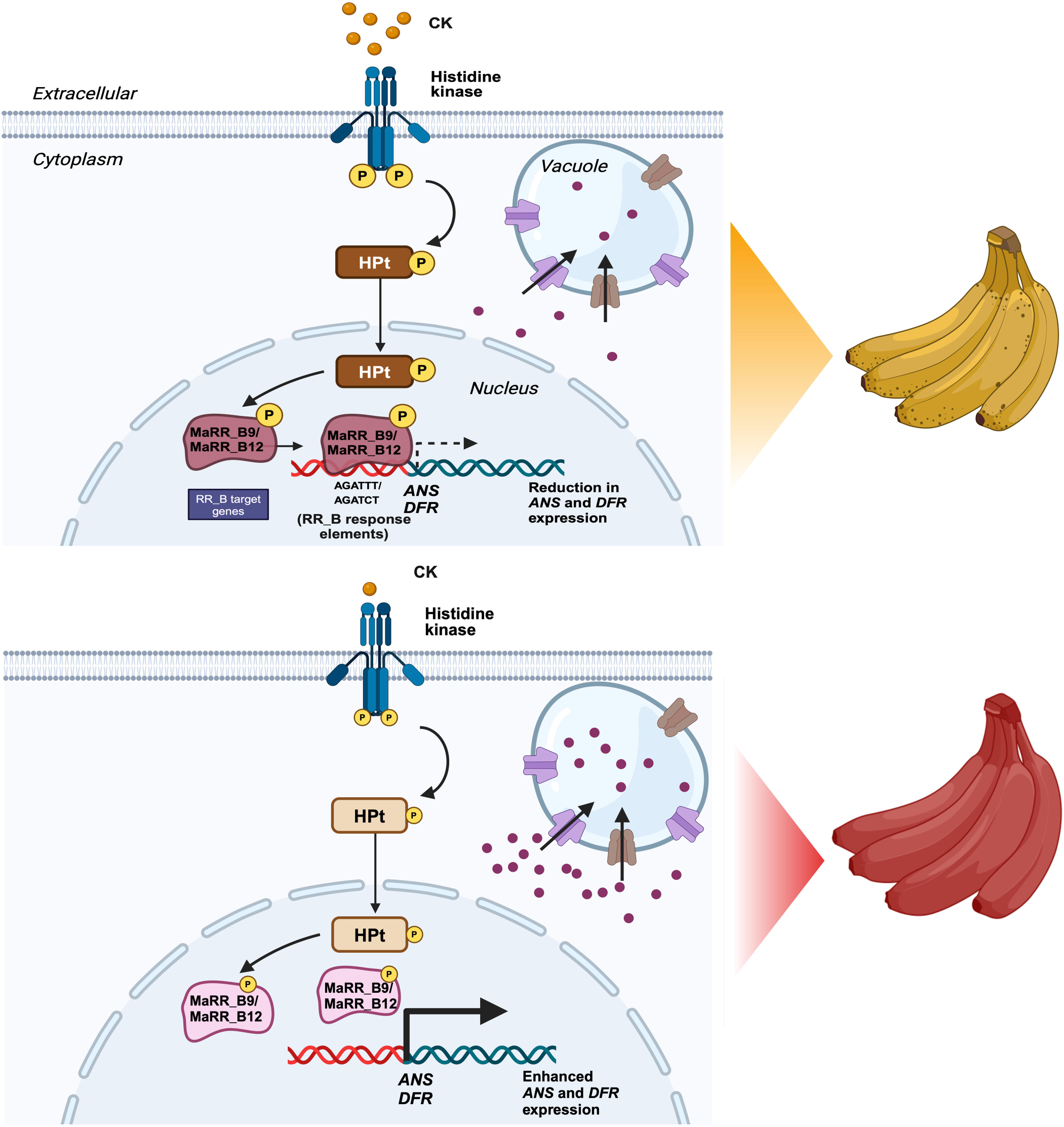
MaRR_B directly binds to the *MaDFR2* and *MaANS* promoters and regulates their expression to modulate AN biosynthesis in banana: (a) Gene structure for the candidate target genes and the primer positions employed used for assessing MaRR_B binding to *proMaDFR* and (b) *proMaANS*. The ATG denotes the initiation codon. The horizontal bars, labeled as fragments a, b, or c, denote the specific target fragments analyzed via chromatin immunoprecipitation-quantitative PCR (ChIP-qPCR). (c) ChIP-qPCR assay showing binding of MaRR_B (AGATnn) to *proMaDFR2* and (d) *proMaANS*. Promoter fragments carrying RR binding elements was co-infiltrated into *N. benthaminana* leaf with *pro35S::MaRR-YFP* construct. Values were normalized with input, which are samples without immunoprecipitation. An anti-IgG antibody was used as a negative control and an anti-GFP antibody was used to precipitate MaRR_B enriched chromatin. The enrichment of anti-IgG antibody was set to 1 and Fold enrichment values are represented as mean ±SD of two biological replicates. Unpaired t-tests were used for statistical analysis. *, p ≤ 0.05; **, p ≤ 0.01, ***, p ≤ 0.001, ns, not significant.

### Transient overexpression of MaRR_B9 and MaRR_B12 decreases anthocyanidin accumulation in unripe banana fruits

To further examine the regulatory role of *MaRR_B9* and *MaRR_B12*, we conducted transient overexpression of these transcription factors in unripe banana fruit slices. Each transcription factor was driven by the maize *Ubiquitin1* (*ZmUBI1*) promoter, while the *GUS* reporter was controlled by the *UBI1* promoter from *Phaseolus vulgaris* (*proPvUBI1*) to serve as the control for transient transformation. The successful transformation of fruit slices was confirmed by the observed GUS activity (Fig. 8a). RT-qPCR analysis corroborated the increased expression of *MaRR_B9* and *MaRR_B12* in *proZmUBI:MaRR_B9* and *B12* overexpressing banana fruits compared to the empty vector (EV) controls (Fig. 8b). However, the relative transcript levels of *MaDFR1*, *MaDFR2*, and *MaANS* were reduced in fruits with transient overexpression of *MaRR_B9* and *MaRR_B12* (Fig 8c).

**Figure 8.**
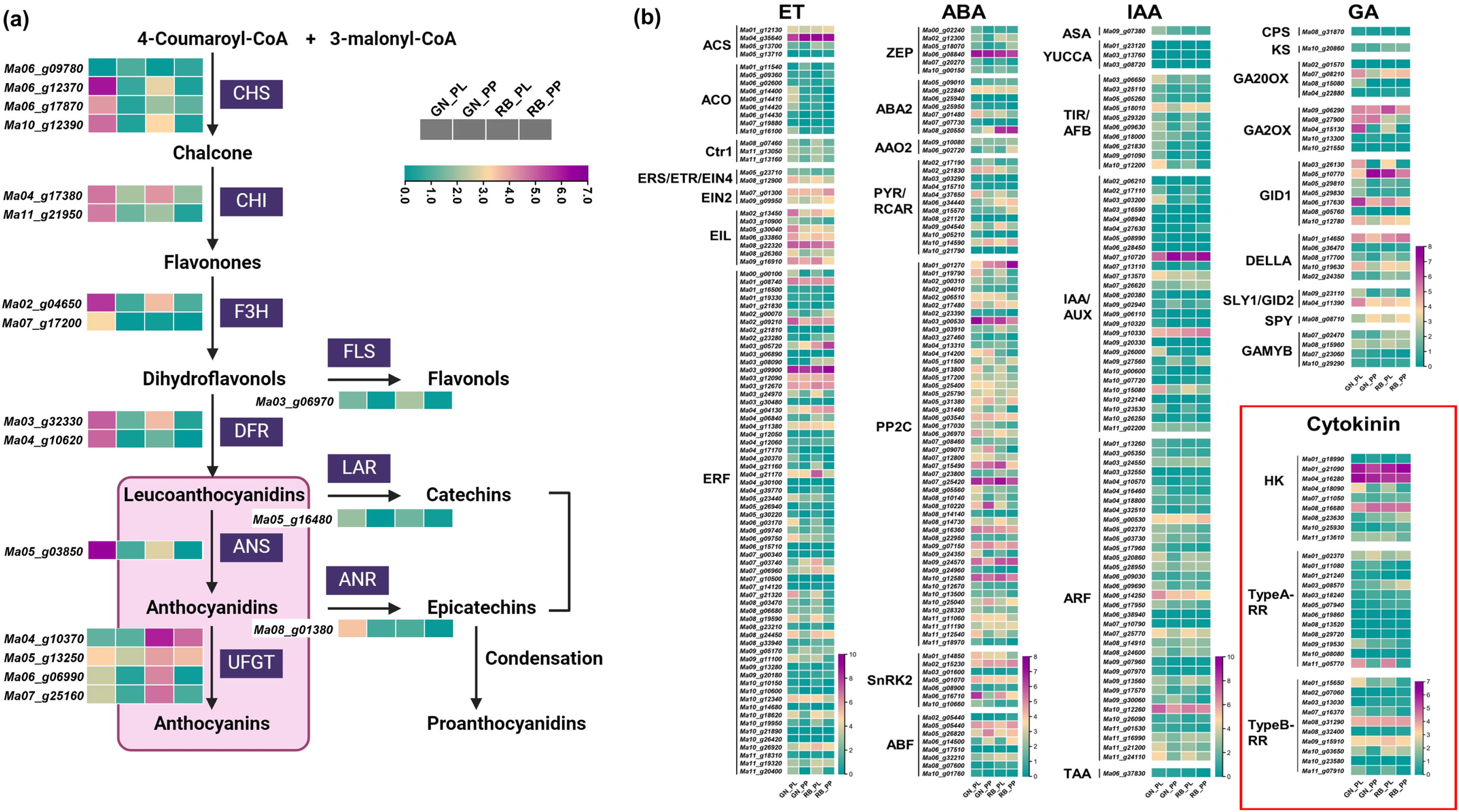
Functional analysis of MaRR_B genes in banana fruits. (a) The *ZmUBI* promoter was used to drive the overexpression of *MaRR_B9* and *MaRR_B12* together with the GUS reporter construct driven by the pea (*Phaseolus vulgaris*) *Ubi1* promoter as an indicator of transient transformation. GUS staining. (b) Cyanidin content (AN-derivative) in transiently transformed fruit slices and control (empty vector). (c) Relative transcript levels of *MaRR_B9*, *MaRR_B12* (d) *MaDFR1*, *MaDFR2* and *MaANS* in transiently transformed fruit slices.

Additionally, we quantified the levels of cyanidin, an anthocyanin derivative, in the transformed fruit slices using LC-MS-based quantification. The results revealed decreased levels of cyanidin in transgenic fruit discs (Fig 8d). In summary, these findings collectively suggest that *MaRR_B9* and *MaRR_B12* negatively regulate anthocyanin biosynthesis by influencing the expression of key anthocyanin biosynthesis genes, specifically *MaDFR* and *MaANS*.

## Discussion

Despite our growing understanding of anthocyanin biosynthesis in different fruit crops, relatively little is known about the fine-tuning regulatory loops governing this crucial metabolic pathway in monocots. The anthocyanin biosynthesis pathway responds to a variety of developmental and environmental cues, indicating the presence of a regulatory cascade involving both positive and negative regulators. Adopting a systematic approach to characterize banana-color variation, we performed a combination of comparative transcriptomics, physiological, molecular, and biochemical studies in the present work. The transcriptome analysis provided valuable insights into the molecular mechanisms underlying the color contrast between these two varieties. These varieties showed enrichment in secondary metabolic processes crucial for their distinct coloration in ripe peel tissue. Since plant hormones have an important role in fruit development and ripening, we focused our study on the relatively understudied area of cytokinin-mediated regulation of anthocyanin biosynthesis in fruit tissues of GN and RB.

Earlier studies suggested the role of phytohormones in the modulation of anthocyanin biosynthesis in different plant species, including fruits. For example, exogenous application of abscisic acid (ABA) and jasmonic acid increased anthocyanin content in fruits like grapevine, apple, sweetcherry (Shen *et al*., 2014; An *et al*., 2015; Sun *et al*., 2019). On the other hand, gibberellins and ethylene negatively regulate anthocyanin accumulation in Arabidopsis and apple (Ni *et al*., 2019; Wang *et al*., 2022). Furthermore, Gibberellic acid and jasmonic acid antagonistically regulate anthocyanin biosynthesis by tight regulation of DELLA and JAZ repressors (Xie *et al*., 2016; An *et al*., 2024). Our transcriptome analysis also revealed altered expression of various hormone-related genes in GN and RB indicating their probable role in anthocyanin biosynthesis. Few studies have also suggested the role of cytokinins (CKs) in anthocyanin biosynthesis (Deikman & Hammer, 1995; Das *et al*., 2012). However, molecular components involved in CK-mediated regulation of anthocyanin biosynthesis remains elusive. Therefore, in the present study, we focused our study on molecular players involved in the CK signaling, as revealed following comparative transcriptome analysis.

Cytokinin signaling in plants occurs through the Two-Component System (TCS) which is involved in responding to various environmental stimuli (Zschiedrich *et al*., 2016; Alvarez & Georgellis, 2023). It typically consists of a histidine kinase (HK), histidine phosphotransfer (HpT) and a response regulator (RR). However, there is a complete lack of knowledge regarding the contribution of type-B RRs to the regulation of anthocyanin biosynthesis in monocots. Type-B RRs have diverse functional roles in TCS signaling and various studies have suggested their role in stress and developmental pathways (Zubo & Schaller, 2020; Yamburenko *et al*., 2020). To this end, genome-wide identification of RRs along with other signaling components has been carried out in the present study. The MaRRs comprising Rec_Myb domain and clustered together with known Type-B RRs from rice and Arabidopsis, were classified as Type-B RRs (Tiwari *et al*., 2021). Phylogenetic clustering and comparative expression profiling indicated that MaRR_B9 and MaRR_B12 might be negative regulators of anthocyanin biosynthesis. Interestingly, CKs have been found to have both negative and positive effects on anthocyanin biosynthesis in various plant species (Das *et al*., 2012; Bhaskar *et al*., 2021; Li *et al*., 2021; Zhu *et al*., 2023). In our study, exogenous application of 30 µM isopentenyladenine (iP) significantly increased the expression of *MaRR_B9* and *MaRR_B12* and reduced expression of two key structural genes of anthocyanin biosynthesis (*DFR* and *ANS,)* and leading to a subsequently reduction in anthocyanin in fruit tissues, which confirmed the negative regulation of anthocyanin biosynthesis in banana by cytokinin. The molecular components of cytokinin signaling are expected to directly mediate this negative regulation of anthocyanin biosynthesis. The upregulated expression of two B-type RRs following cytokinin treatment strengthened our hypothesis that MaRR_B9 and MaRR_B12 are directly involved in the cytokinin mediated regulation of anthocyanin.

The type-B *Arabidopsis* RRs are the positive regulators in the two-component cytokinin signaling pathway. The triple mutants *arr1-3 arr10-5 arr12-1* of type-B RRs display altered anthocyanin content in Arabidopsis (Argyros *et al*., 2008). In another study, Das *et al*. (2012) demonstrated that cytokinin regulation of anthocyanin biosynthesis is channeled primarily through type-B RRs. Given this, the present investigation suggests that cytokinin type-B RRs of the TCS might be involved in the cytokinin-mediated differential regulation of anthocyanin biosynthesis in GN and RB, two color contrasting varieties of banana.

B-ARR binds to specific *cis-*acting elements present in the promoter regions of their target genes. Previously, B-ARR binding sites were identified in the wuschel promoter by which it directs shoot development in Arabidopsis (Xie *et al*., 2018). B-type ORRs could also bind to the promoter of type-A ORR genes with the B-RR binding sites on their promoters (Shi *et al*. 2020). We also identified putative B-RR binding motifs in the promoters of *MaDFR* and *MaANS*, suggesting that candidate B-RRs (MaRR_B9 and MaRR_B12) might directly regulate their transcription and thereby could modulate anthocyanin pathway. Other than B-ARR motif, in our previous study, we also identified other *cis*-regulatory elements for abscisic acid response element (ABRE) sites and low-temperature response in these promoters (Rajput *et al*., 2022a) which suggests a possible role of different regulators in modulating the AN pathway. For example, FaRAV1, an AbA-responsive TF, regulates anthocyanin biosynthesis by interacting with and increasing the expression of the *CHS*, *F3H*, *DFR* and *GT1* promoters (Zhang *et al*., 2020). In another example, light responsive TFs PIF3 and HY5 activate the expression of anthocyanin-specific biosynthesis genes by binding to their light responsive units and thereby positively regulating anthocyanin biosynthesis (Shin *et al*., 2007).

We further demonstrated that MaRR_B9 and MaRR_B12 interact with the promoters of *MaDFR* and *MaANS* and negatively modulate their activity. Furthermore, overexpression of *MaRR_B9* and *MaRR_B12* genes in banana fruits also downregulates anthocyanin biosynthesis, resulting in reduced anthocyanin accumulation. However, in Arabidopsis, cytokinin treatment induces anthocyanin biosynthesis by involving type-B RR (Das *et al*., 2012). These observations suggest that CK might modulate anthocyanin biosynthesis differentially in different plant species. (Zhu *et al*., 2023). These reports also highlight lineage or species-specific regulation of anthocyanin biosynthesis by growth regulators. In various studies, type-B RRs were shown to act as both positive and negative regulators. For example, rice *rr21/22/23* mutant supported both negative and positive roles for type-B RRs in regulating root growth based on its concentration (Worthen *et al*., 2019). Also, Lineage-specific expansions of the type-B and type-A RR gene families have occurred in both monocots and dicots (Pils & Heyl, 2009; Tsai *et al*., 2012). This implies that while these gene families share common ancestral functions, monocot RRs have likely acquired additional, distinct functions beyond those observed in dicots (Worthen *et al*., 2019). We also found in our previous study the lack of correlation between flavonoid-related metabolites and transcript levels in pulp tissue. Our findings hinted that some additional upstream or parallel regulators in addition to components of MBW complex, might be involved in the regulation of flavonoid biosynthesis (Rajput *et al*., 2022a). These type-B RRs might be the potential upstream regulators that suppress the expression of biosynthesis genes.

Based on our findings, we propose a model illustrating how cytokinin mediates the regulation of anthocyanin biosynthesis through MaRR_B9 and MaRR_B12. Upon cytokinin induction, these proteins get phosphorylated and bind to the promoters of anthocyanin biosynthetic genes, such as ANS and DFR, thereby reducing their expression (Fig. 9). This study advanced our knowledge of the molecular basis of anthocyanin biosynthesis in monocots, in which cytokinin mediated regulation is less explored than in dicots and offers targets involved in anthocyanin biosynthesis for genome editing for banana biofortification. The findings of the present study deepen our understanding of the complex signaling pathways and highlights the broader implications of cytokinin signaling in anthocyanin biosynthesis. Further research in this direction may uncover additional components of this regulatory network, providing the way for advancements in banana improvement strategies and the development of other crop plants with higher nutritional value.

**Figure 9.**
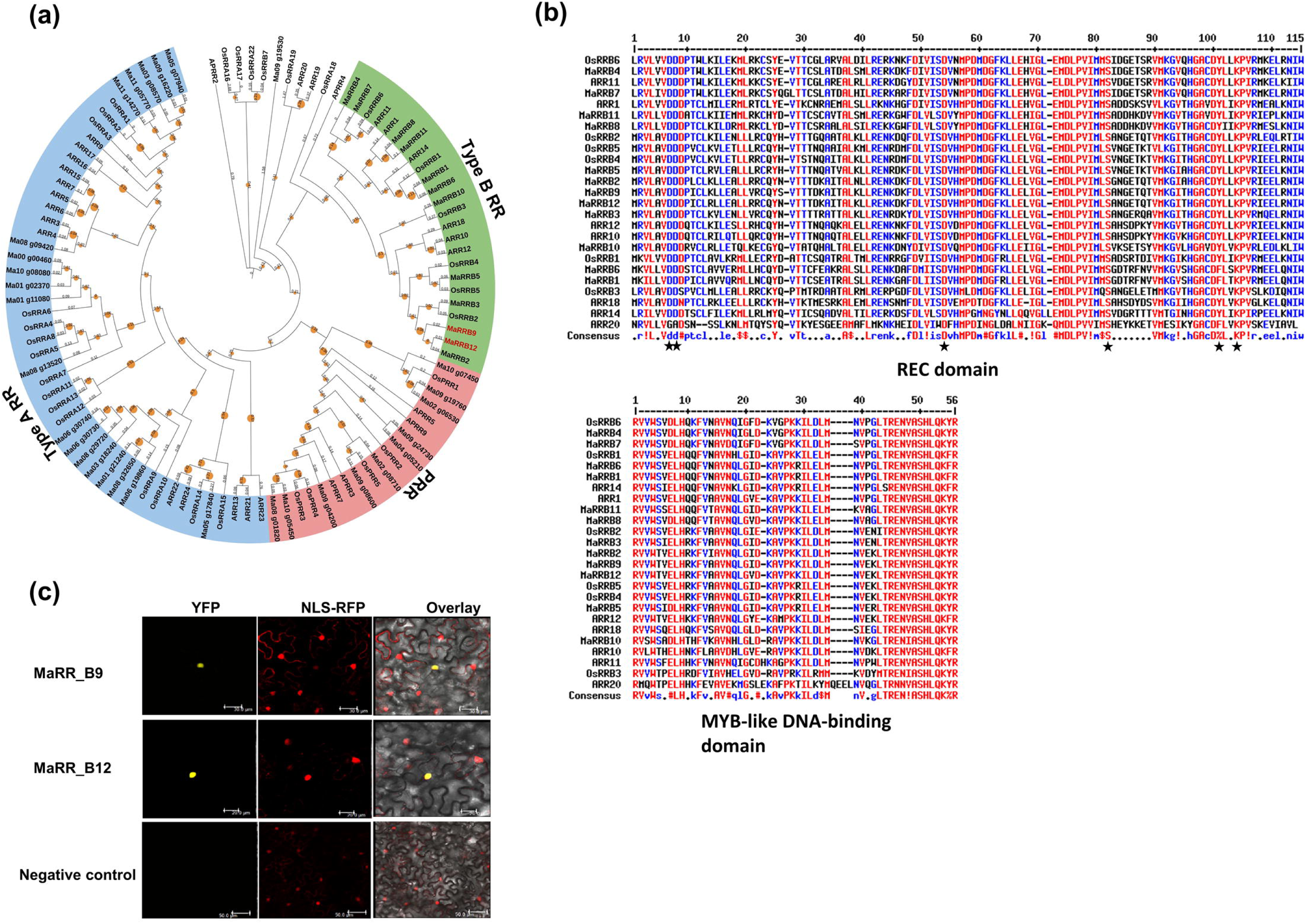
Working model of cytokinin signaling pathway in Grand Naine and Red Banana. Upon cytokinin induction, Histidine Kinases (HK) phosphorylate Histidine Phosphotransfer proteins (HPTs), which in turn transfer the phosphoryl group to response regulators, MaRR_B9 and MaRR_B12. In Grand Naine banana, phosphorylated MaRR_B9 and MaRR_B12 directly bind to the B-RR sites of target gene promoters of anthocyanin biosynthetic enzymes including *ANS* and *DFR*, leading to reduced expression of *ANS* and *DFR* (marked by fine black arrow), resulting in reduced anthocyanin content and less transport of anthocyanin from the cytosol to the vacuole. Conversely, in red banana, lower cytokinin concentrations lead to reduced phosphorylation of HKs, HPTs, MaRR_B9, and MaRR_B12. This reduces the interaction of MaRR_B9 and MaRR_B12 with the *MaANS* and *MaDFR* promoters, resulting in increased expression of *MaANS* and *MaDFR* (marked by solid black arrow) and consequently, increased anthocyanin content and more transport of anthocyanin from the cytosol to the vacuole.

## Acknowledgements

This work was supported by the core grant of National Institute of Plant Genome Research and Department of Science and Technology-SERB for Core Research Grant to AP. RR acknowledges Council of Scientific and Industrial Research, Government of India, for Senior Research Fellowship. ST acknowledges the Department of Biotechnology, for a Research Associate (RA) Fellowship. The authors are thankful to DBT-eLibrary Consortium (DeLCON) for providing access to e-resources. We acknowledge the Metabolome facility at NIPGR for phytochemical analysis.

## Author’s contribution

RR, ST, and AP conceived the idea and designed the research. RR, ST, and KA conducted experiments. RR, ST, and AP interpreted the data. AL & PM provides insightful suggestions during discussion. RR, ST, PM, and AP wrote the manuscript. AP coordinated the research project. All authors read and approved the final manuscript.

## Data availability

All data supporting the findings of this study are available within the paper and within the Supplementary data published online.

## Declarations

The authors declare no conflict of interest.

## Method S1 Functional annotation and pathway enrichment analysis

The DEGs obtained in different combinations were utilized for gene ontology (GO) mapping and enrichment analysis by using agriGO v2.0 platform to determine their involvement in specific biological processes or signaling pathways. Statistically significant GO terms (significance level ≤0.05) obtained after applying Fisher statistical method and Yekutieli adjustment method were considered for the GO enrichment analysis. The KEGG pathway enrichment analysis was performed for DEGs using DAVID functional annotation tool at p-Value <0.05 (Sherman *et al*., 2022). To conduct pathway analysis in different tissues, firstly we created a MapMan mapping file for *M. acuminata* gene model sequences using the Mercator tool. This file categorizes genes into bins based on hierarchical ontologies obtained from various databases. We then used MapMan v.3.6.0RC1 (Thimm *et al*., 2004) to visualize the differentially expressed genes (DEGs) in GN_PL, GN_PP, RB_PL and RB_PP.

## Supplementary Information

**Supplementary figure 1. Gene ontology and enrichment analysis of differentially expressed genes (DEGs).** (a) Gene ontology (GO) analysis of up-regulated and down-regulated DEGs showing enrichment in biological processes, (b) cellular components, and (c) molecular functions. The significantly enriched GO terms are represented in the bar graph, with the y-axis showing the -log10(p-value) of enrichment and the x-axis indicating the GO terms.

**Supplementary figure 2. KEGG pathway enrichment analysis of differentially expressed genes (DEGs).** (a) Enriched KEGG pathways for up-regulated DEGs, and (b) down-regulated DEGs. Enrichment analysis was performed using the DAVID functional annotation tool at p-Value <0.05.

**Supplementary figure 3. Mapman Analysis of Differentially Expressed Genes (DEGs).** Mapman analysis showing the functional overview of DEGs in (a) Grand naine (GN) peel; (b) red banana (RB) peel; (c) GN pulp; (d) GN pulp. The analysis highlights the regulation overview represented by different bins. Upregulated genes are depicted in red, while downregulated genes are shown in blue. The size of the bins corresponds to the number of genes associated with each functional category.

**Supplementary figure 4. Multiple sequence alignment of type B response regulators (RRs).** The MSA of type B response regulators (RRs) from *A. thaliana*, *O. sativa*, and *M. acuminata*. The MSA was generated using full-length protein sequences and highlights the conserved REC domain and MYB-like DNA binding domain among these species.

**Supplementary figure 5. Phylogenetic analysis and multiple sequence alignment of histidine kinases.** (a) A phylogenetic tree was constructed using the full-length protein sequences of histidine kinases from *A. thaliana*, *O. sativa*, and *M. acuminata*. The tree shows clustering into three groups, labeled as I, II, and III, which are highlighted in different shades. The tree was generated using the maximum-likelihood method at 1000 bootstrap values. (b) The Multiple Sequence Alignment (MSA) was performed using the multAlin and highlights the conserved CHASE domain, His KA domain, and REC domain.

**Supplementary figure 6. Phylogenetic analysis and multiple sequence alignment of histidine phosphotransferase.** (a) A phylogenetic tree was generated based on the full-length protein sequences of histidine phosphotransferase (HpTs) from *A. thaliana*, *O. sativa*, and *M. acuminata*. The tree displayed clustering into I, II, and III groups highlighted in varying shades. The tree was constructed using the maximum-likelihood method with 1000 bootstrap replications. (b) Multiple Sequence Alignment (MSA) of MaHpTs, AtHpTs and OsHpTs was conducted using multAlin. Highly conserved residues across these species are highlighted in red.

**Supplementary figure 7. Expression analysis of candidate Histidine Kinase and type-A Response regulatory genes from *M. acuminata.*** (a). RT-qPCR expression analysis of *MaHK* and (b) *MaRR_A* genes were carried out in GN_PL vs RB_PL and GN_PP vs RB_PP tissues. The graphs show values ±SD of three technical replicates. *MaActin* was used as reference control.

**Supplementary figure 8. Expression analysis of candidate type-B Response regulatory genes from *M. acuminata.*** RT-qPCR expression analysis of *MaRR_B* genes were carried out in GN_PL vs RB_PL and GN_PP vs RB_PP tissues. The graphs show values ±SD of three technical replicates. *MaActin* was used as reference control.

**Supplementary figure 9.** Representation of *MaDFR1* promoter with one putative *cis*-acting RR site (AGATAT) shown with yellow triangle. Transcription start site (TSS) is indicated with − 1 position.

**Supplementary file S1.** Comprehensive lists of differentially expressed genes (DEGs) identified in ripe peel (PL) and pulp (PP) tissue comparisons within Grand naine (GN) and Red banana (RB) cultivars. It includes up- and downregulated genes in GN_PL vs GN_PP, GN_PL vs RB_PL, GN_PP vs RB_PP, and RB_PL vs RB_PP. These DEGs were identified based on their log2(fold_change) values, calculated at significant p-values and q-values. Additionally, the FPKM values for different tissues and annotations for the corresponding proteins are mentioned.

**Supplementary file S2.** Gene ontology (GO) mapping for different comparisons of tissues at significant p-value and FDR. The GO terms are categorized into three main categories: biological process (P), molecular function (F), and cellular component (C).

**Supplementary file S3.** KEGG pathway analysis of up- and down-regulated DEGs in different combinations.

**Table S1.** List of primers used in the present study.

**Table S2.** Read alignment summary.

