## Supplementary Information for "Type-B response regulators MaRR_B9 and MaRR_B12 coordinate cytokinin-mediated negative regulation of anthocyanin biosynthesis in banana fruits"

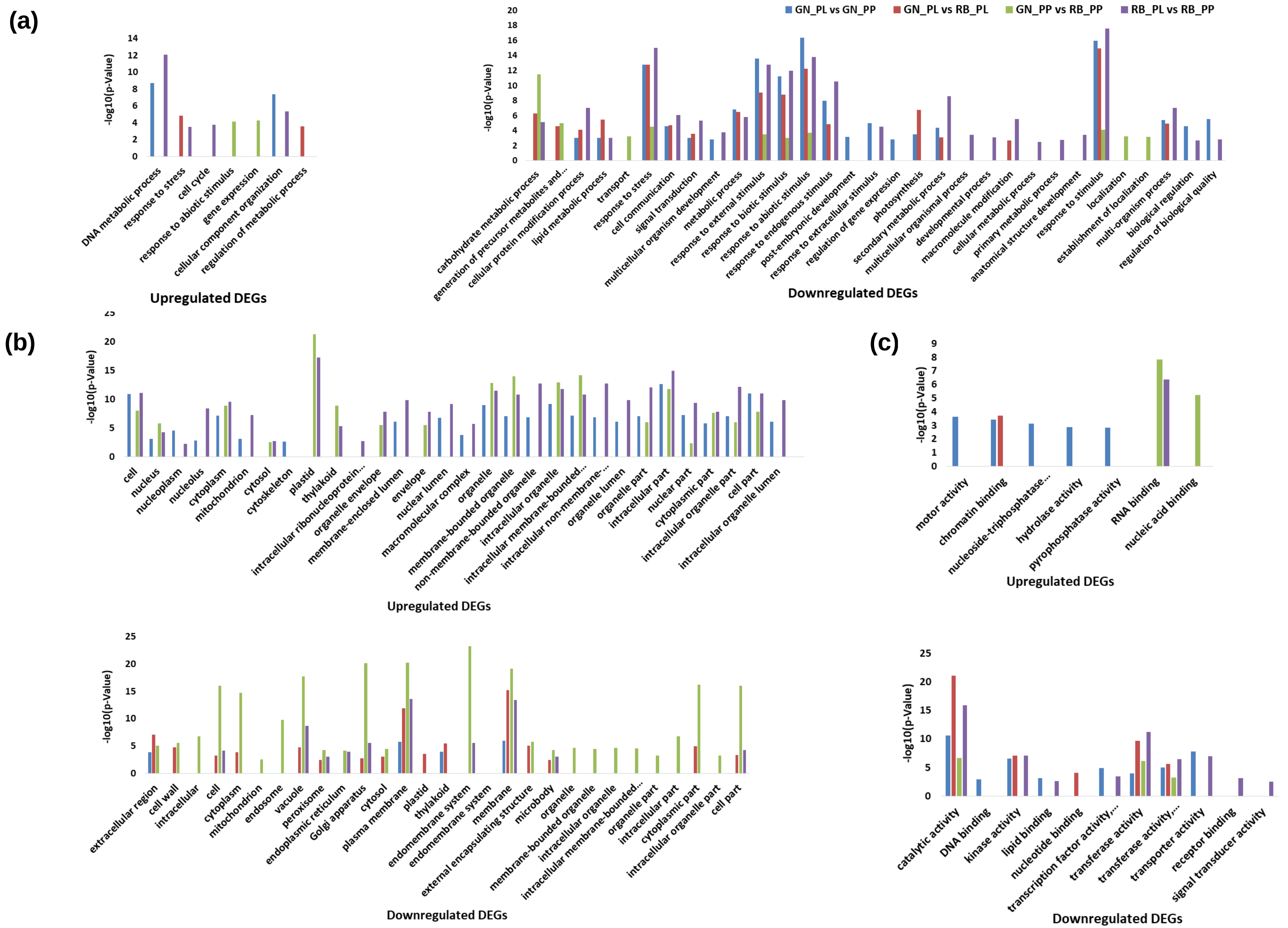
(a)
-log10(p-Value)
DNA metabolic process
response to stress
cell cycle
response to abiotic stimulus
gene expression
cellular component organization
regulation of metabolic process
Upregulated DEGs
GN_PL vs GN_PP
GN_PL vs RB_PL
GN_PP vs RB_PP
RB_PL vs RB_PP
carbohydrate metabolic process
generation of precursor metabolites and...
cellular protein modification process
lipid metabolic process
transport
response to stress
cell communication
signal transduction
multicellular organism development
metabolic process
response to external stimulus
response to biotic stimulus
response to abiotic stimulus
response to endogenous stimulus
post-embryonic development
response to extracellular stimulus
regulation of gene expression
photosynthesis
secondary metabolic process
multicellular organismal process
developmental process
macromolecule modification
cellular metabolic process
primary metabolic process
anatomical structure development
response to stimulus
localization
establishment of localization
multi-organism process
biological regulation
regulation of biological quality
Downregulated DEGs
(b)
cell
nucleus
nucleoplasm
nucleolus
cytoplasm
mitochondrion
cytosol
cytoskeleton
plastid
thylakoid
intracellular ribonucleoprotein...
organelle envelope
membrane-enclosed lumen
envelope
nuclear lumen
macromolecular complex
organelle
membrane-bounded organelle
non-membrane-bounded organelle
intracellular organelle
intracellular membrane-bounded...
intracellular non-membrane-...
organelle lumen
organelle part
intracellular part
nuclear part
cytoplasmic part
intracellular organelle part
cell part
intracellular organelle lumen
Upregulated DEGs
extracellular region
cell wall
intracellular
cell
cytoplasm
mitochondrion
endosome
vacuole
peroxisome
endoplasmic reticulum
Golgi apparatus
cytosol
plasma membrane
plastid
thylakoid
endomembrane system
endomembrane system
membrane
external encapsulating structure
microbody
organelle
membrane-bounded organelle
intracellular organelle
intracellular membrane-bounded...
organelle part
intracellular part
cytoplasmic part
intracellular organelle part
cell part
Downregulated DEGs
(c)
motor activity
chromatin binding
nucleoside-triphosphatase...
hydrolase activity
pyrophosphatase activity
RNA binding
nucleic acid binding
Upregulated DEGs
catalytic activity
DNA binding
kinase activity
lipid binding
nucleotide binding
transcription factor activity,...
transferase activity
transferase activity,...
transporter activity
receptor binding
signal transducer activity
Downregulated DEGs

Fig. S1

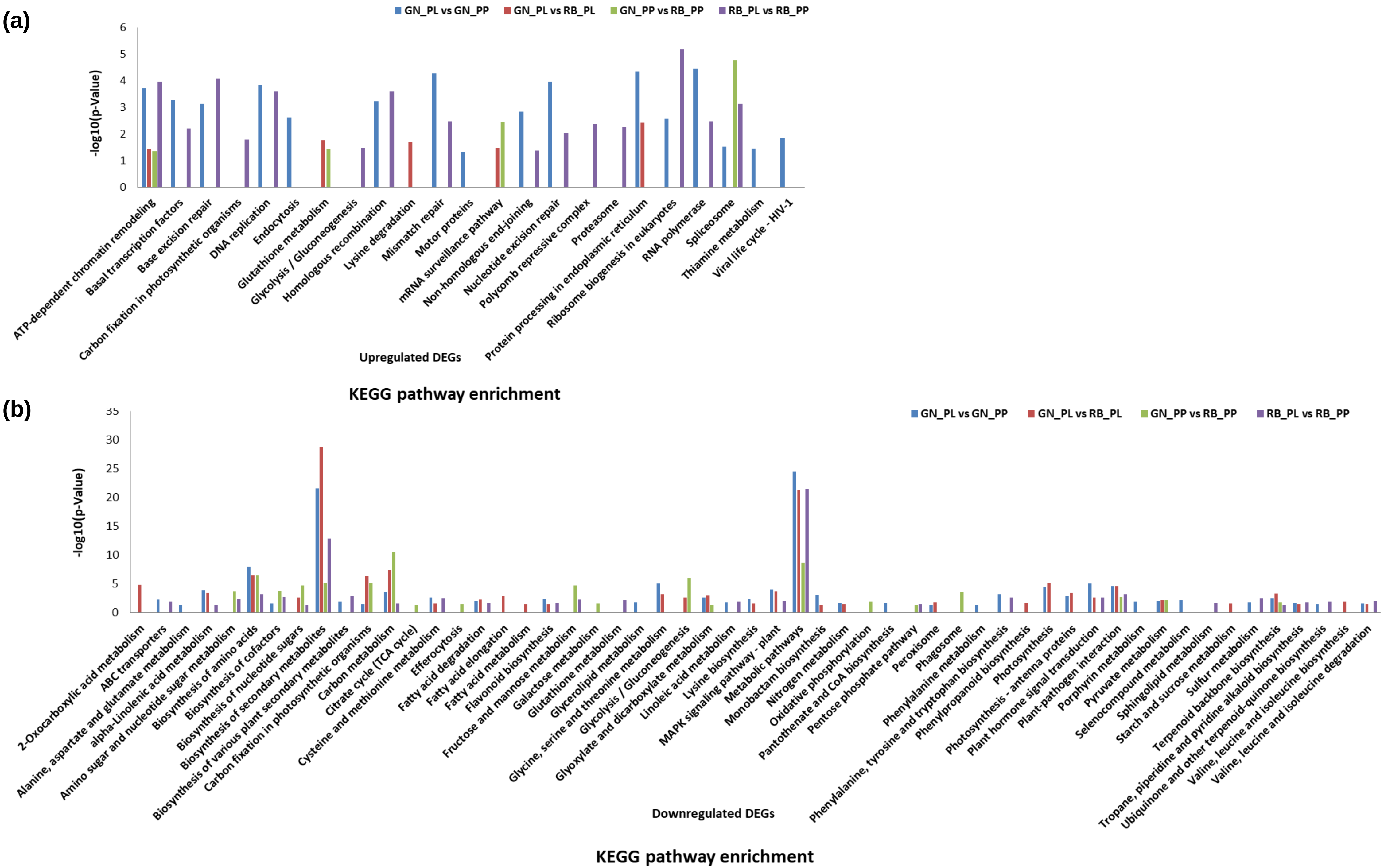
(a)
GN_PL vs GN_PP
GN_PL vs RB_PL
GN_PP vs RB_PP
RB_PL vs RB_PP
-log10(p-Value)
0
1
2
3
4
5
6
ATP-dependent chromatin remodeling
Basal transcription factors
Base excision repair
Carbon fixation in photosynthetic organisms
DNA replication
Endocytosis
Glutathione metabolism
Glycolysis / Gluconeogenesis
Homologous recombination
Lysine degradation
Mismatch repair
Motor proteins
mRNA surveillance pathway
Non-homologous end-joining
Nucleotide excision repair
Polycomb repressive complex
Proteasome
Protein processing in endoplasmic reticulum
Ribosome biogenesis in eukaryotes
RNA polymerase
Spliceosome
Thiamine metabolism
Viral life cycle - HIV-1
Upregulated DEGs
KEGG pathway enrichment
(b)
GN_PL vs GN_PP
GN_PL vs RB_PL
GN_PP vs RB_PP
RB_PL vs RB_PP
-log10(p-Value)
0
5
10
15
20
25
30
35
2-Oxocarboxylic acid metabolism
ABC transporters
Alanine, aspartate and glutamate metabolism
alpha-Linolenic acid metabolism
Amino sugar and nucleotide sugar metabolism
Biosynthesis of amino acids
Biosynthesis of cofactors
Biosynthesis of nucleotide sugars
Biosynthesis of secondary metabolites
Biosynthesis of various plant secondary metabolites
Carbon fixation in photosynthetic organisms
Carbon metabolism
Citrate cycle (TCA cycle)
Cysteine and methionine metabolism
Efferocytosis
Fatty acid degradation
Fatty acid elongation
Fatty acid metabolism
Flavonoid biosynthesis
Fructose and mannose metabolism
Galactose metabolism
Glutathione metabolism
Glycerolipid metabolism
Glycine, serine and threonine metabolism
Glycolysis / Gluconeogenesis
Glyoxylate and dicarboxylate metabolism
Linoleic acid metabolism
Lysine biosynthesis
MAPK signaling pathway - plant
Metabolic pathways
Monobactam biosynthesis
Nitrogen metabolism
Oxidative phosphorylation
Pantothenate and CoA biosynthesis
Pentose phosphate pathway
Peroxisome
Phagosome
Phenylalanine metabolism
Phenylalanine, tyrosine and tryptophan biosynthesis
Phenylpropanoid biosynthesis
Photosynthesis
Photosynthesis - antenna proteins
Plant hormone signal transduction
Plant-pathogen interaction
Porphyrin metabolism
Pyruvate metabolism
Selenocompound metabolism
Sphingolipid metabolism
Starch and sucrose metabolism
Sulfur metabolism
Terpenoid backbone biosynthesis
Tropane, piperidine and pyridine alkaloid biosynthesis
Ubiquinone and other terpenoid-quinone biosynthesis
Valine, leucine and isoleucine biosynthesis
Valine, leucine and isoleucine degradation
Downregulated DEGs
KEGG pathway enrichment

Fig. S2

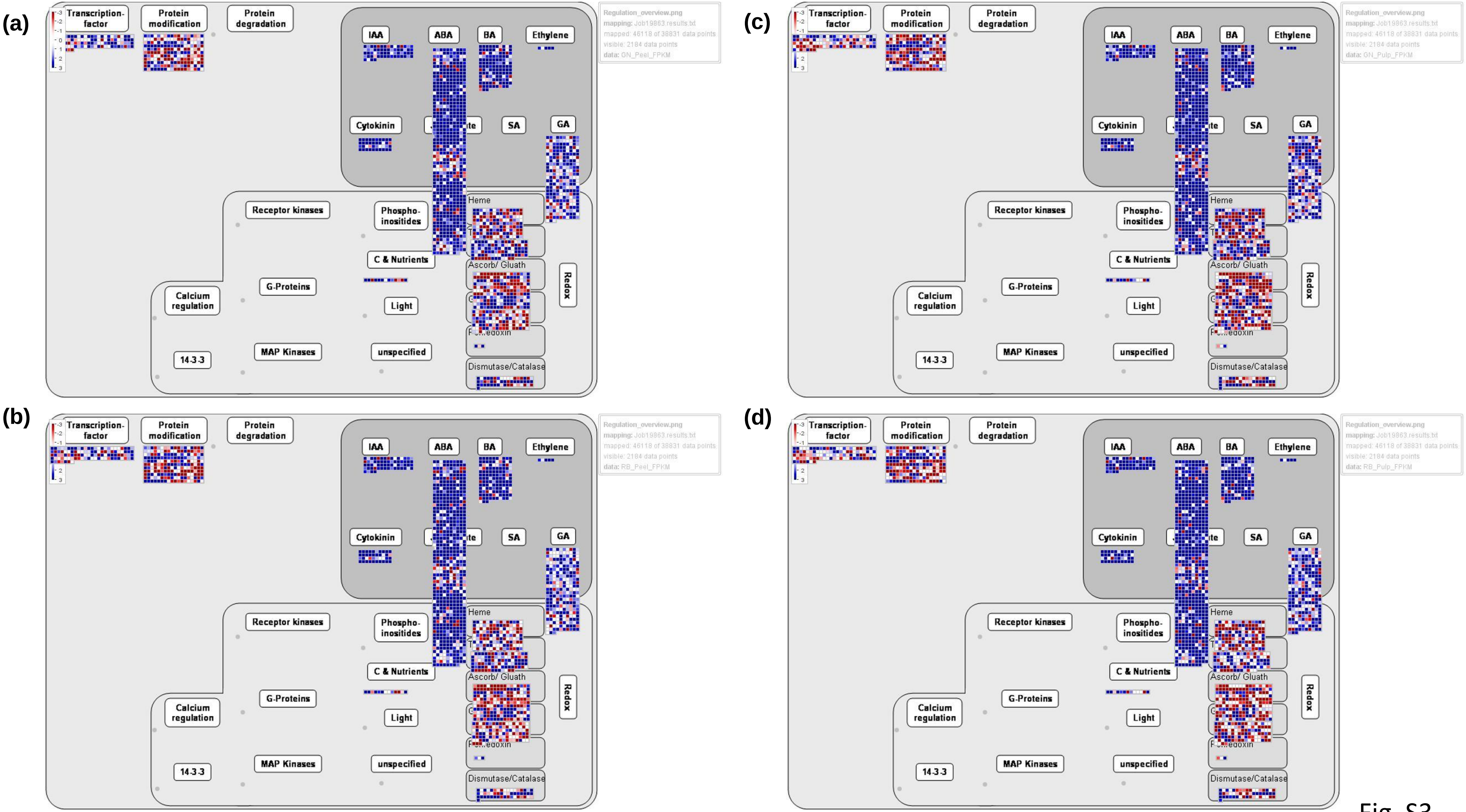
(a)
(b)
(c)
(d)
Transcription-factor
Protein modification
Protein degradation
IAA
ABA
BA
Ethylene
Cytokinin
SA
GA
Receptor kinases
Phospho-inositides
C & Nutrients
G-Proteins
Light
Calcium regulation
14-3-3
MAP Kinases
unspecified
Heme
Ascorb/ Gluath
Dismutase/Catalase
Redox
Regulation_overview.png
mapping: Job19863.results.txt
mapped: 46118 of 38831 data points
visible: 2184 data points
data: GN_Peel_FPKM
data: GN_Pulp_FPKM
data: RB_Peel_FPKM
data: RB_Pulp_FPKM

Fig. S3

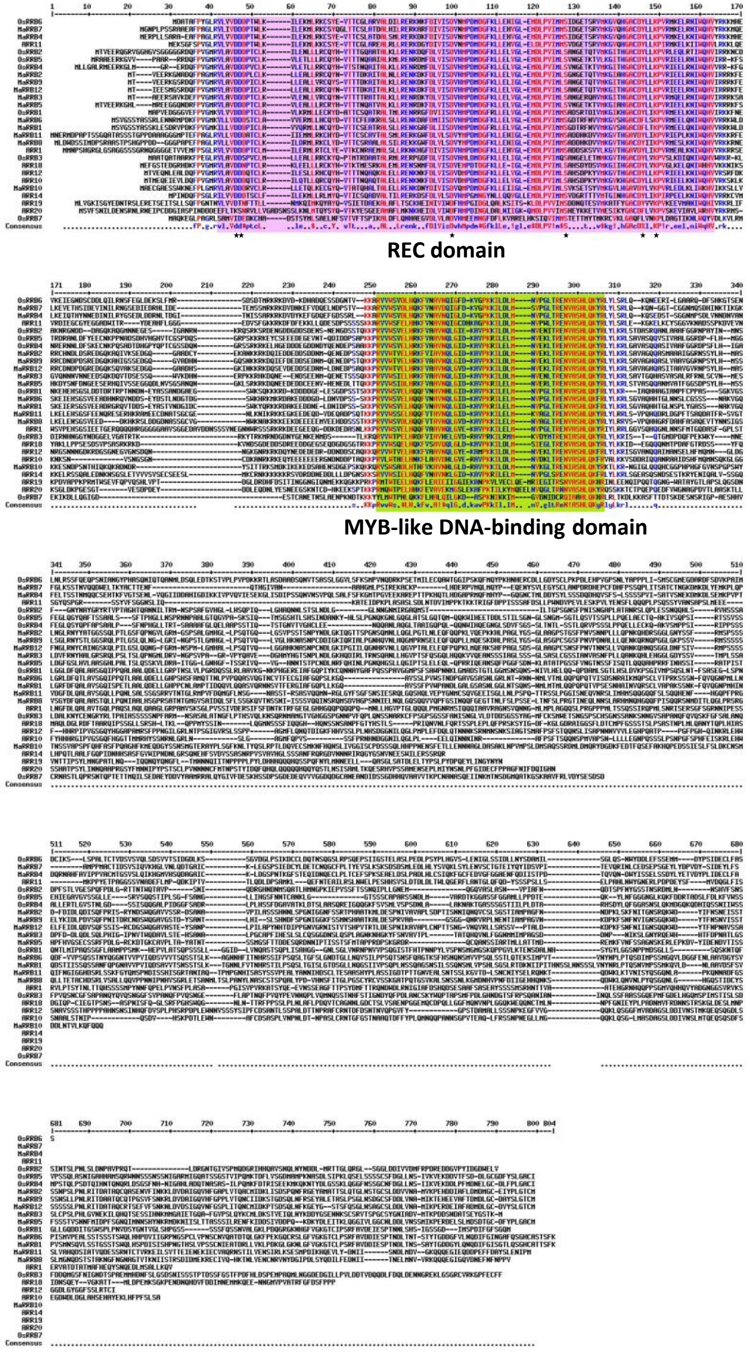


Fig. S4

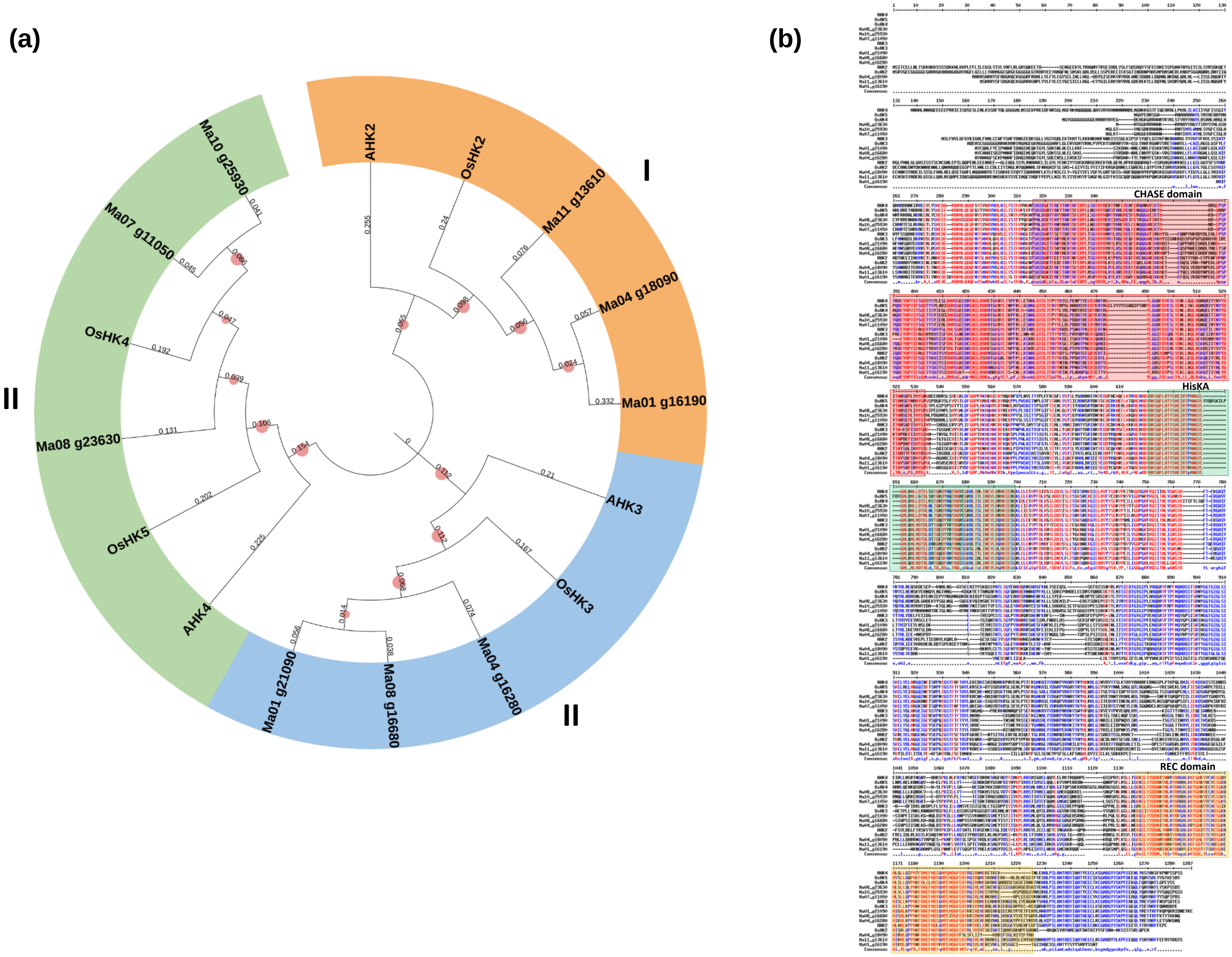
(a)
(b)
I
II
III
AHK2
OsHK2
Ma11 g13610
Ma04 g18090
Ma01 g16190
AHK3
OsHK3
Ma04 g16280
Ma08 g16680
Ma01 g21090
AHK4
OsHK5
Ma08 g23630
OsHK4
Ma07 g11050
Ma10 g25930
CHASE domain
HisKA
REC domain

Fig. S5

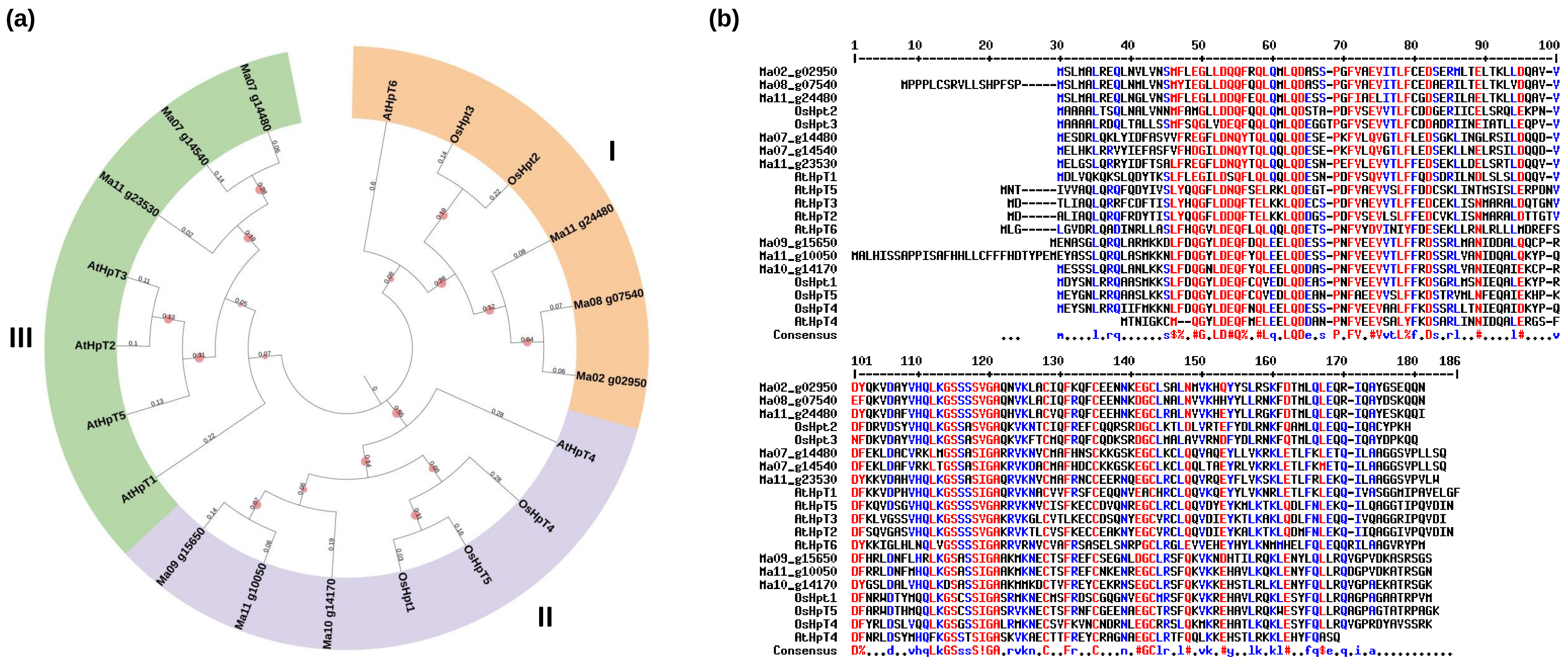
(a)
(b)
I
II
III
AtHpT6
OsHpt3
OsHpt2
Ma11 g24480
Ma08 g07540
Ma02 g02950
AtHpT4
OsHpT4
OsHpT5
OsHpt1
Ma10 g14170
Ma11 g10050
Ma09 g15650
AtHpT1
AtHpT5
AtHpT2
AtHpT3
Ma11 g23530
Ma07 g14540
Ma07 g14480
Ma02_g02950
Ma08_g07540
Ma11_g24480
OsHpt2
OsHpt3
Ma07_g14480
Ma07_g14540
Ma11_g23530
AtHpT1
AtHpT5
AtHpT3
AtHpT2
AtHpT6
Ma09_g15650
Ma11_g10050
Ma10_g14170
OsHpt1
OsHpT5
OsHpT4
AtHpT4
Consensus

Fig. S6

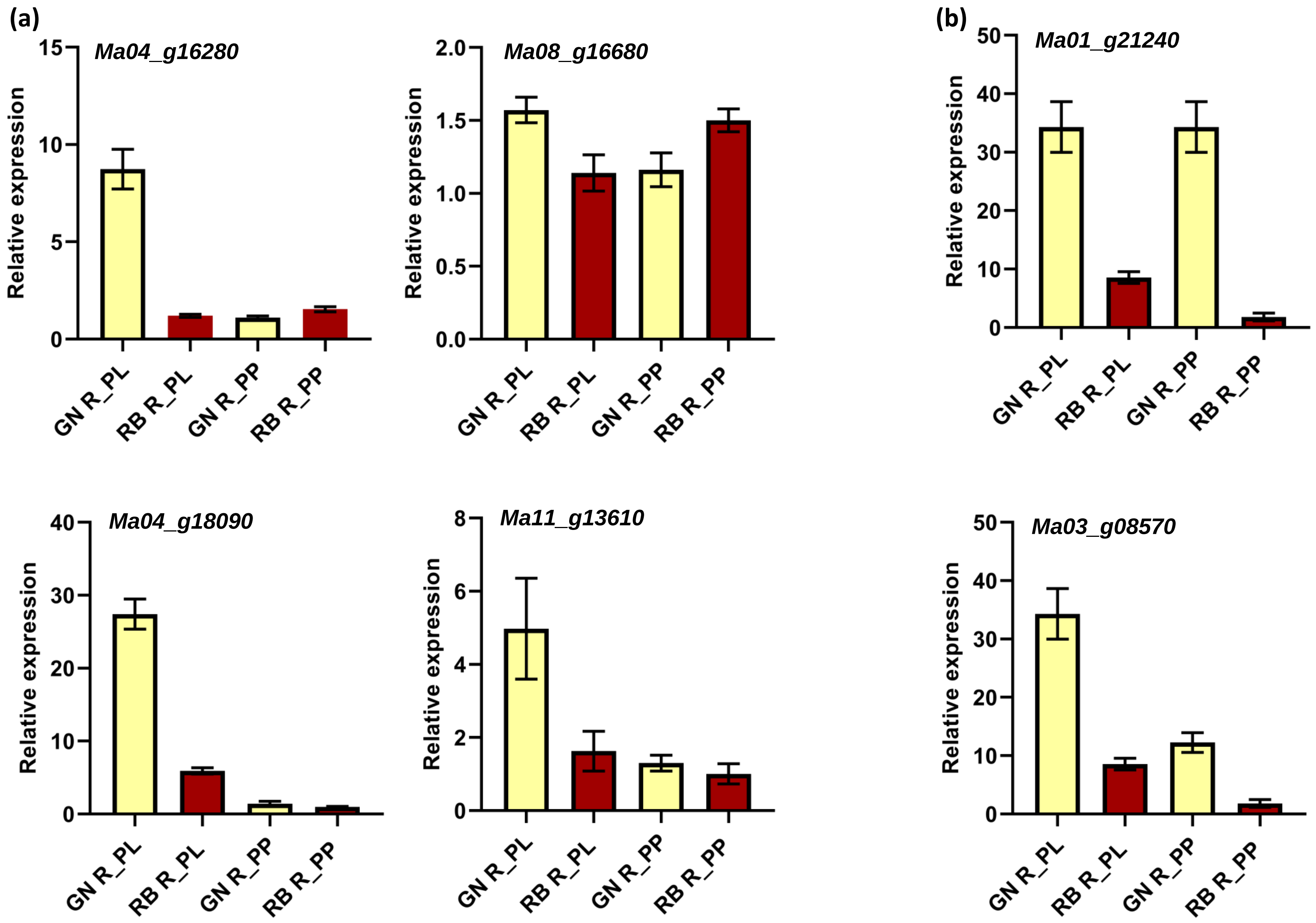

(a)
Ma04_g16280
Ma08_g16680
Ma04_g18090
Ma11_g13610
(b)
Ma01_g21240
Ma03_g08570
Relative expression
GN R_PL
RB R_PL
GN R_PP
RB R_PP


Fig. S7

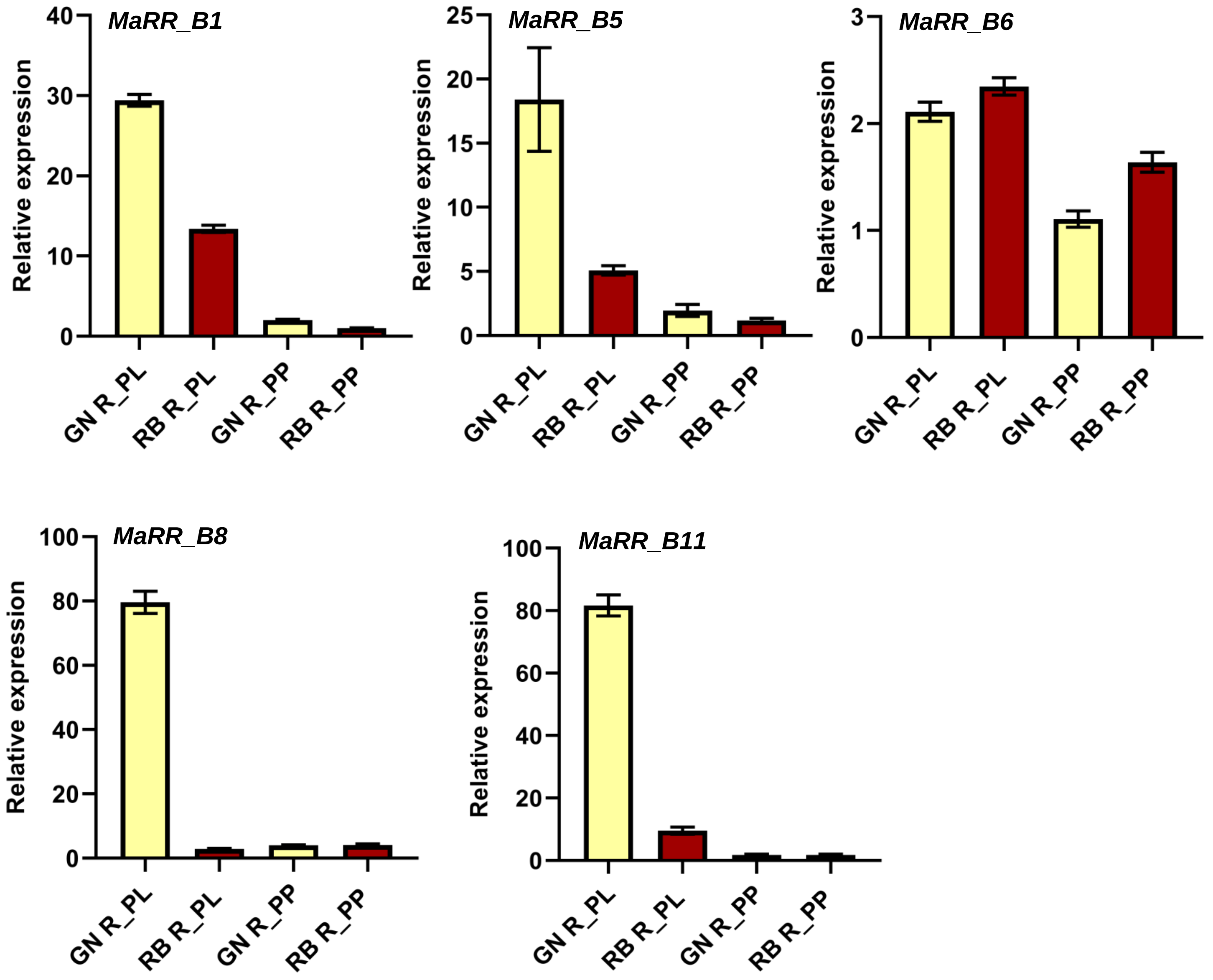
MaRR_B1
MaRR_B5
MaRR_B6
MaRR_B8
MaRR_B11
Relative expression
GN R_PL
RB R_PL
GN R_PP
RB R_PP
0
10
20
30
40
5
15
25
1
2
3
60
80
100

Fig. S8

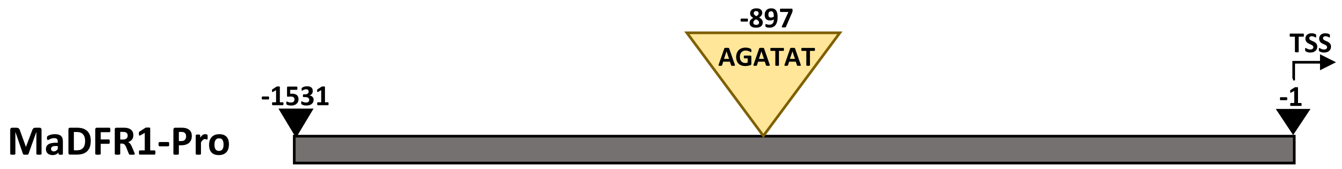


Fig. S9

**Table S1.** List of primers used in the present study.

| Primer Name | Sequence (5′ to 3′) | Purpose |
|---|---|---|
| MaANS_For | 5′GAGGGGAAGCTGGACAGGGAAC3′ | RT-qPCR forward primer |
| MaANS_Rev | 5′GTTGTGGAGGATGAAGGAGAGC3′ | RT-qPCR reverse primer |
| MaDFR1_For | 5′CGAACGGGTGCACTTCTCCTCC3′ | RT-qPCR forward primer |
| MaDFR1_Rev | 5′CACTGCACTGTTCCTCTGCTGT3′ | RT-qPCR reverse primer |
| MaDFR2_For | 5′CACAGGTTTCTTCTTGACCATCTCCAG3′ | RT-qPCR forward primer |
| MaDFR2_Rev | 5′AGACATCTTAAGGCTGCACTTCTCC3′ | RT-qPCR reverse primer |
| Ma04_g16280_For | 5′GACTGCTTTTGAGAGGCCACTGACG3′ | RT-qPCR forward primer |
| Ma04_g16280_Rev | 5′GTGCGGGTGTCTGCTCTGTTGGAT3′ | RT-qPCR reverse primer |
| Ma08_g16680_For | 5′CGTTGACCAGTGGTGTAGCATATG3′ | RT-qPCR forward primer |
| Ma08_g16680_Rev | 5′TCCTTGCACACGGGATGTCTCTTC3′ | RT-qPCR reverse primer |
| Ma04_g18090_For | 5′GCTAGGATGTCATGGAGAAGGG3′ | RT-qPCR forward primer |
| Ma04_g18090_Rev | 5′GTCGTCCAATCCCCATAATAACC3′ | RT-qPCR reverse primer |
| Ma11_g13610_For | 5′CCGAAGGTGGAGCAACCCGTTG3′ | RT-qPCR forward primer |
| Ma11_g13610_Rev | 5′CGTCTTCTCTCTCTCATCTCCCAA3′ | RT-qPCR reverse primer |
| Ma01_g21240_For | 5′AGGAGCAGGTGAAGAAAGCCGGG3′ | RT-qPCR forward primer |
| Ma01_g21240_Rev | 5′ACAACCCTCCACTGTTCTTGAGCAAC3′ | RT-qPCR reverse primer |
| Ma03_g08570_For | 5′GCTGTTGTTGCAGAAGCCCAATTCCA3′ | RT-qPCR forward primer |
| Ma03_g08570_Rev | 5′CAAGAGCCCTGCTCCCAGAATCC3′ | RT-qPCR reverse primer |
| MaRR_B1_For | 5′GTACAGGTAGCAGCACTAGTTGGCA3′ | RT-qPCR forward primer |
| MaRR_B1_Rev | 5′CCGAGATTTTTCCCTAGTCCCTTGTC3′ | RT-qPCR reverse primer |
| MaRR_B5_For | 5′GACAGTTGAGGAGAGGAAGGGGC3′ | RT-qPCR forward primer |
| MaRR_B5_Rev | 5′CGAAGCAGCAGAGCCTCTAACACC3′ | RT-qPCR reverse primer |
| MaRR_B6_For | 5′TGGGACAAGTCTTAGATGGAGGCTTTG3′ | RT-qPCR forward primer |
| MaRR_B6_Rev | 5′AGGCCGACCTATAATAACATCAGGGTG3′ | RT-qPCR reverse primer |
| MaRR_B8_For | 5′GGATCCCAGCAGAGCAGCGTC3′ | RT-qPCR forward primer |
| MaRR_B8_Rev | 5′GCGGAGCATCCTGTCGAGGATCTT3′ | RT-qPCR reverse primer |
| MaRR_B9_For | 5′CAGACACCAGGAAGTGTCTTTTCCAAC3′ | RT-qPCR forward primer |
| MaRR_B9_Rev | 5′CCGTCTCCAGTGCATATTCACTCC3′ | RT-qPCR reverse primer |
| MaRR_B11_For | 5′GCCAAGCCACCGCCAGTAGCT3′ | RT-qPCR forward primer |
| MaRR_B11_Rev | 5′ACTTGCGGAGCATCATCTCGATGATC3′ | RT-qPCR reverse primer |
| MaRR_B12_For | 5′CTGCAAATCCCTTGCCTATACTTGCTC3′ | RT-qPCR forward primer |
| MaRR_B12_Rev | 5′GTTGGATTACAAAGAGGTGCAACAGCC3′ | RT-qPCR reverse primer |
| MaRR_B9_For | 5′GGGGACAAGTTTGTACAAAAAAGCAGGCTccATGACG GTGGAGGAGAGGAAGGG3′ | attB1 forward primer for full length cloning of MaRR_B9 cDNA into entry clone |
| MaRR_B9_Rev | 5′GGGGACCACTTTGTACAAGAAAGCTGGGTaTCACATG CAAGTGCCTAAAGAGTAG3′ | attB2 reverse primer for full length cloning of MaRR_B9 cDNA into entry clone |

| | | |
|---|---|---|
| MaRR_B12_For | 5′GGGGACAAGTTTGTACAAAAAAGCAGGCTccATGACGATTGAAGAGAGCATGGG3′ | attB1 forward primer for full length cloning of MaRR_B12 cDNA into entry clone |
| MaRR_B12_Rev | 5′GGGGACCACTTTGTACAAGAAAGCTGGGTaTCACATGCAAGTACCTAAAGAGTAG3′ | attB2 reverse primer for full length cloning of MaRR_B12 cDNA into entry clone |
| proMaDFR2_For | 5′GGGGACAAGTTTGTACAAAAAAGCAGGCTccGATTAAGCCAGCTCATCTACTAGT3′ | attB1 forward primer for cloning of MaDFR2 promoter into entry clone for dual luciferase assay |
| proMaDFR2_Rev | 5′GGGGACCACTTTGTACAAGAAAGCTGGGTaTTTAAAGACTGAAACCAGAGGGAG3′ | attB2 reverse primer for cloning of MaDFR2 promoter into entry clone for dual luciferase assay |
| proMaANS_For | 5′GGGGACAAGTTTGTACAAAAAAGCAGGCTccTTTATACTCTTAGCACTACGATTTTGGG3′ | attB1 forward primer for cloning of MaANS promoter into entry clone for dual luciferase assay |
| proMaANS_Rev | 5′GGGGACCACTTTGTACAAGAAAGCTGGGTaTAGGAGAAGCTTCCGCTACTGGCAGCAA3′ | attB2 reverse primer for cloning of MaANS promoter into entry clone for dual luciferase assay |
| MaDFR _1_For | 5′TCATACCGCAACATATATTCATTAGC3′ | CHIP-qPCR forward primer |
| MaDFR_1_Rev | 5′CCATTCAAATTAGAATGTTATTTCCTTG3′ | CHIP-qPCR reverse primer |
| MaDFR_2_For | 5′CTAGTCAAATTCTAAATCATTCAGAAT3′ | CHIP-qPCR forward primer |
| MaDFR _2_Rev | 5′TTACTGTTTATCTTTAATCTTATCTGG3′ | CHIP-qPCR reverse primer |
| MaDFR _3_For | 5′TTCATATCAACAAACAAGGAAGTAG3′ | CHIP-qPCR forward primer |
| MaDFR_3_Rev | 5′CGGATCGTAATCACGTCAAATAT3′ | CHIP-qPCR reverse primer |
| MaANS_1_For | 5′GGTGCGCACGTGCTTCCCTTT3′ | CHIP-qPCR forward primer |
| MaANS_1_Rev | 5′CGTGGGGCCACGCTCTACAAA3′ | CHIP-qPCR reverse primer |

**Table S2. Read alignment summary**

| Sample Name | Total Read Count | Read Count after rRNA removal | QC Pass % | Aligned Read Count | Aligned % | Unaligned % |
|---|---|---|---|---|---|---|
| GN Peel_R1 | 61,240,560 | 60,493,974 | 98.78% | 57,592,695 | 95.20% | 4.80% |
| GN Peel_R2 | 78,844,290 | 71,434,902 | 90.60% | 68,461,233 | 95.84% | 4.16% |
| GN Peel_R3 | 75,740,324 | 73,280,618 | 96.75% | 70,411,617 | 96.08% | 3.92% |
| GN pulp_R1 | 87,902,178 | 79,234,442 | 90.14% | 76,534,387 | 96.59% | 3.41% |
| GN pulp_R2 | 87,336,306 | 79,652,900 | 91.20% | 76,701,321 | 96.29% | 3.71% |
| GN pulp_R3 | 98,211,028 | 90,016,562 | 91.66% | 86,402,333 | 95.98% | 4.02% |
| RB peel_R1 | 68,112,612 | 65,635,978 | 96.36% | 62,701,674 | 95.53% | 4.47% |
| RB peel_R2 | 79,421,854 | 78,790,016 | 99.20% | 75,562,375 | 95.90% | 4.10% |
| RB peel_R3 | 66,198,110 | 65,809,452 | 99.41% | 63,269,738 | 96.14% | 3.86% |
| RB pulp_R1 | 81,692,762 | 81,465,628 | 99.72% | 78,508,681 | 96.37% | 3.63% |
| RB pulp_R2 | 65,678,952 | 65,474,154 | 99.69% | 63,064,062 | 96.32% | 3.68% |
| RB pulp_R3 | 57,791,172 | 57,567,046 | 99.61% | 55,466,824 | 96.35% | 3.65% |
